# Naturalistic behavior and self-generated neural activity predictive of self-correction

**DOI:** 10.64898/2026.05.26.727951

**Authors:** Shijie Gu, Chenyan Liu, Anna K. Gillespie, Rhino Nevers, Eric L. Denovellis, Michael E. Coulter, Kenneth Kay, Loren M. Frank

## Abstract

Changing one’s mind –– revising a past decision independently of external cues –– can lead to better outcomes, but its neural basis remains poorly understood. We therefore developed a challenging spatial task for rats where a conflict between innate foraging biases and task rules leads to abundant, spontaneous, and characteristically corrective changes-of-mind (COMs). Neural recordings in the hippocampus, a brain region implicated in counterfactual thinking, revealed two distinct stages wherein more local representations gave way to representations of distant alternatives after animals had begun to reverse course. These representations predicted their eventual choice, and often serially encoded trial start and end locations. Our novel task paradigm reveals distinct representational phases engaged during self-correction and uncovers a rich repertoire of hippocampal spatial representations tied to behavior.

## Main Text

Revising a decision retrospectively and independently of external cues –– i.e., performing a “change-of-mind” (COM) –– can be beneficial for optimal decision-making, despite incurring costs such as energy and time (*1, 2*). COM behavior has been observed across various animal species, including mice, rats, macaques, and humans (*1, 3–8*). In the brain, COM implies one or multiple internal processes that monitor the current decision, determine that it is not preferred, and then redirect decisions toward alternative options. Prior work aimed at understanding the neural substrates of COM relied on ambiguous sensory stimuli (*3, 6, 9, 10*) or cued responses (*5, 9, 11*), making it difficult to isolate internally-driven processes, and most studies included only binary alternatives, limiting focus to error monitoring (*1, 3, 5, 6, 9–12*). As a result, how the various processes engaged during COM unfold over time and across brain regions (*1, 3, 11, 8, 13–15*) remains poorly understood.

To address these challenges, we developed a novel cognitive task – specifically for freely behaving rats in a maze environment – that exploits a conflict between the rats’ innate bias and externally provided task rules. This yielded abundant discrete COM events — rats entering one arm, stopping, reversing, turning and traveling to another. When combined with neural recordings, this behavioral paradigm enabled us to probe neural activity patterns underlying COM and subsequent (revised) decisions. We found evidence for a specific role for population-level activity in the hippocampus, a brain region linked to planning, on-line memory-guided decision-making, and counterfactual thinking (*16–21*).

### Rat spatial sequence task and change-of-mind behavior

We designed a task requiring navigation and permitting free behavior to promote ethological responses, in both behavior and neural activity. Our task had three components: (i) more than two discrete alternatives, (ii) a requirement to learn a challenging rule that is implicit (i.e., does not rely on a direct external cues for a given navigational choice), and (iii) a conflict between innate (preexisting) spatial alternation tendencies/biases and task reward contingencies (*22*). Beyond broadly enabling ethological behavioral responses, we hypothesized that components (ii) and (iii) could lead to naturalistic COM responses that could engage deliberative processes (*11, 13, 14*).

The resulting task, which we call the spatial sequence task (Figure 1A) takes place in a maze consisting of a small central platform connected to a home arm (0.4 m) on one side, and four choice arms (1 m each) on the other (Figure S1A). The end of each arm contains a reward well with a photobeam that dispenses liquid reward (evaporated milk + 5% sucrose) upon a nose-poke if the well was visited in the correct sequence. In a pre-training period, rats learned to start each trial by poking at the home well, proceed to finish each trial by nose-poking at one of the outer arm wells, and then return to the home well to start another trial. Following pretraining, rewards were delivered based on an un-cued spatial sequence (Figure 1A): rewards in a given choice arm (e.g. arm 4) only became available after the animal had obtained reward at a specific previous arm (e.g. arm 1) and then returned to the home well. Thus, a correct sequence might be: home – 1 – home – 4 – home – 2 – home – 3 – home – 1. Erroneous nose-pokes in an outer arm reward well did not reset the sequence and animals were required to return and nose-poke at the home well before initiating their next outer arm choice. Importantly, rats could revise their initial arm choice simply by reversing course and exiting the arm before choosing another arm.

**Figure 1.**
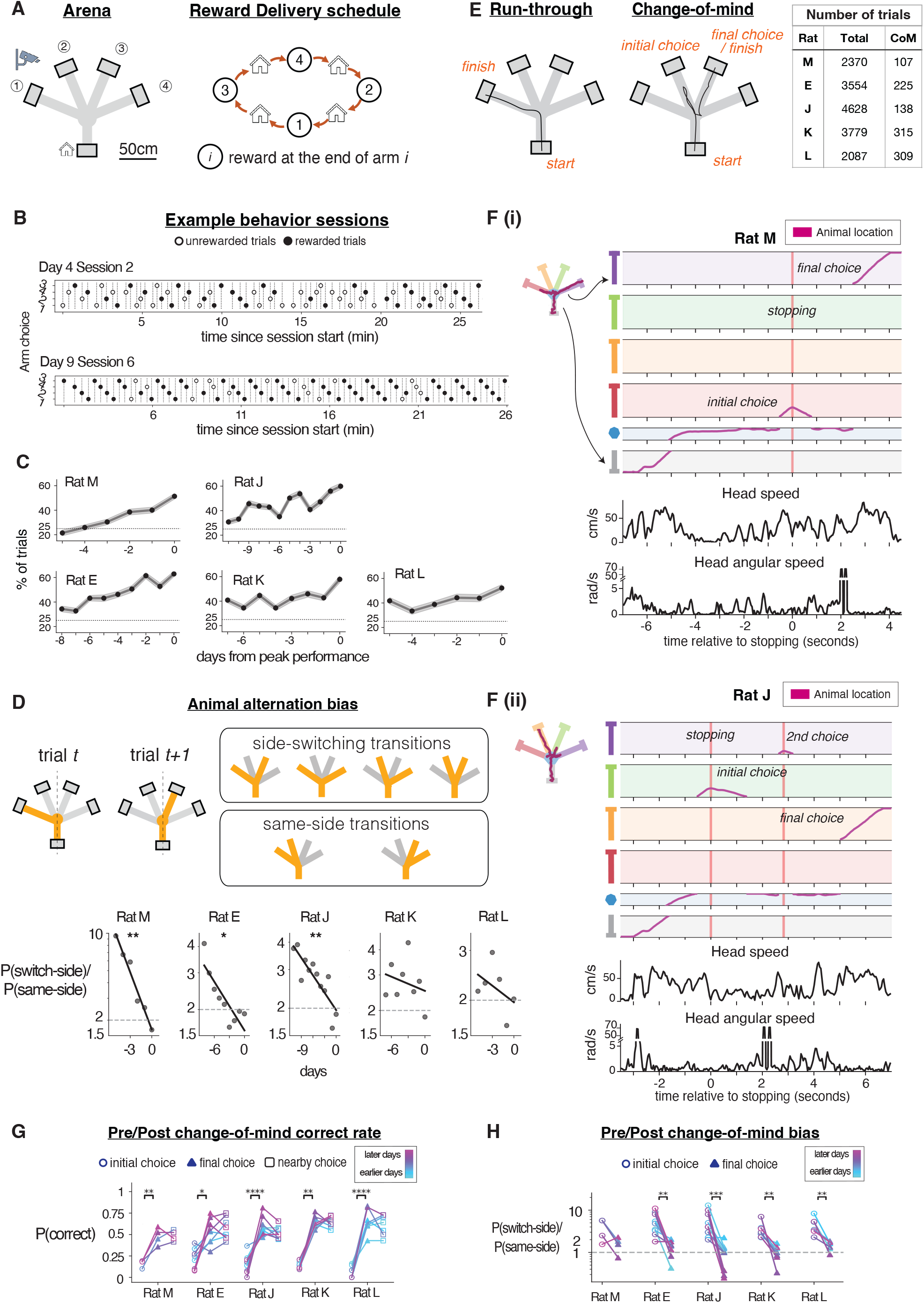
Spatial sequence task, task performance, and change-of-mind behavior. A. Left: schematic of maze and task, consisting of 4 outer choice arms and a home arm, each ending in a reward well. Each trial is initiated by a rat subject poking at the well at the home arm and ends with a choice in an outer choice arm. Right: schematic illustrating the order of the reward delivery schedule among the reward wells. B. Example snippets of choices of the subject Rat E over time in one session. Arm choices are ordered by row from top to bottom in the correct reward sequence (3-4-2-1-3). Unrewarded trials are denoted by hollow circles; rewarded trials are denoted by solid circles. C. Learning curves showing the percentage of consecutive trials that conform to correct sequence transitions over time as rats learned the task. D. Top: Schematic illustrating side-switching (subject chose between consecutive arms that are on opposite sides of the maze) and same-side transitions (subject chose between consecutive arms that are on the same side of the maze). Bottom: Ratio of probabilities of side-switching vs same-side transitions across days. E. Schematic illustrating change-of-mind (COM) trials. The subject would enter one choice arm before poking in another choice arm reward port. Inset: the total number of trials recorded in this dataset and the number of COM trials per subject. F. Example change-of-mind trials with animal trajectories. The top panel shows the 1D projections of the 2D animal location in the maze (shown in the top left) in magenta. The maze segments are separated and organized as shown by colors in the top left. Lower panels show animal 2D head speed as well as angular speed. (i) Rat M. (ii) Rat J. In example (ii), rat J leaves the home well first, then enters arm 3 (shaded green in the plot) from the central platform before stopping (marked by the red vertical bar). Rat J then explores a bit of arm 4 (shaded purple), before making its final choice to arm 2 (shaded orange). G. Daily average probability of obtaining a reward at the initial choice (hollow circles) on a COM trial, the final choice (solid triangles) on a COM trial and on a temporally nearby run-through trials with no COMs (hollow squares). Each data point represents an aggregation from a single day. In all figures in this paper:*p<0.05, **p<0.01, ***p<0.001, ****p<0.0001. H. Daily ratio of probabilities of subjects making switch-side choices and same-side choices during the initial choice (denoted by hollow circles) and final choice (denoted by solid triangles) on COM trials. Each data point is data aggregated from a single day. Stars label significant differences from paired t-tests, p < 0.01 for all rats except M. In addition, sign-test for comparison between the initial ratio and 2 yields p < 0.01 for rat J and K, p < 0.05 for rat E and L, between the final ratio and 2 yields p < 0.05 for rat J, K, E and L, and between final ratio and 1 yield insignificant difference in all rats.

The sequence of reward delivery was designed to invoke a conflict between strategies.Previous studies have documented preexisting alternation strategies in rats to visit less-recently visited places or foraging patches (*22, 23*). Consistent with this, our pilot behavior experiments (not shown) indicated that rats tended to alternate between the left and right overall areas (sides) of the maze, which we term “switch-side” bias (Figure 1B). We therefore chose an order of reward delivery that included a conflict between spatial alternation tendencies and reward contingencies. The resulting sequences have a 1:1 ratio of same-side vs switch-side transitions (sequences 1-2-4-3-1 in 2 rats and 3-4-2-1-3 in 3 rats) requiring animals to overcome their “switch-side” bias. Importantly, this bias both promotes exploration and simultaneously conflicts with completion of the correct reward sequence.

All rats learned the task over a period of 5-19 days (Figure 1B,C, Figure S1E). This improvement in performance was associated with a shift in the prevalence of switch-side vs. same-side transitions (Figure 1D). Initially animals were biased strongly to switch-side transitions (Figure 1D; ratios were greater than 2 for all animals, reflecting the two switch-side transitions available for each trial as compared to the single same-side transition for a random choice strategy; binomial test, p < 0.032). As learning progressed, this ratio decreased to ∼1.5 (mixed generalized logistic effect, mean multiplicative estimate of number of days spent in the maze in reducing bias is 0.92, p < 1e-4).

Surprisingly, this change in transition behavior was associated with frequent COM events. In standard “run-through” trials the rats would enter an outer arm and then proceed directly to the reward port at the end (Figure 1E, left). By contrast, in other trials, the rats fully entered an outer arm, reflecting a decision, but then came to a sudden stop. They then backed up before turning around and choosing a different arm (Figure 1E, right, 1F(a),1F(b), and Figure S1D). We define such behavior as a COM event and a trial with at least one COM event as a COM trial. In total, we identified hundreds of COM trials per subject (Figure 1E inset; see Figure S1E for the prevalence across days). In most COM trials, rats changed their minds once (entered 2 choice arms), though trials with multiple COM events (>2 arms entered in a trial) were also seen in all rats (Figure 1F(b), Figure S1F).

COM trials were predominantly corrective: the final decision in COM trials resulted in higher reward rates than would have been the case had the animal continued with their initial choice (Figure 1G, ranksum tests, individually for each rat, p < 0.01 for rats M, J, K, L and p < 0.05 for rat E). These corrections brought the average performance on COM trials up to the level seen on run-through trials (Figure 1G, no significant differences between COM trials and run-through trials), suggesting that COM is engaged when the animal’s internal monitoring processes detect a likely error.

These errors were associated with corrections for the natural side-switching bias. The initial choice on COM trials was heavily biased to be a side-switch, while the final choice was much less likely to be a side-switch with a ratio of switching to same-side transitions close to 1 (Figure 1H). Thus, our behavioral paradigm elicits many COMs that spontaneously occur, are predominantly corrective, and are consistent with a conflict between learned and innate behavior.

### Hippocampal representations of alternatives are only expressed after stopping

The hippocampus is implicated in planning, memory-guided decision-making (*16, 22, 24, 25*), and counterfactual thinking (*18, 26*) — all relevant to COM. We therefore investigated whether hippocampal activity contributed to the initial decision to stop or the subsequent corrective choice. Rats were implanted with 32-or 64-tetrode microdrive arrays targeting the dorsal hippocampus (Figure S1B, S1C), and a clusterless Bayesian method (*27*) was used to assess the population-level spatial representation present in the hippocampus throughout COM events (Figure S2).

We observed two patterns of representations of alternatives: (1) an lack of spatial representation far from the animal expressed prior to stopping, i.e. before rats came to a stop in the middle of the choice arm, and (2) extensive representation away from the animal once rats stopped, and while they were reversing course toward the center of the maze (Figure 2, S3, Movie S1-S3). These latter representations included spatially remote arms, including the home arm, as well as the reward well region in the current arm (Figure S3). Unlike a previous report of remote representations associated with COM events during immobility(*8*), the non-local activity we observed occurred without animals’ extended movement pauses and occurred exclusively during the hippocampal theta rhythm, which is generally associated with movement (Figure S7, S8, S9, S10).

**Figure 2.**
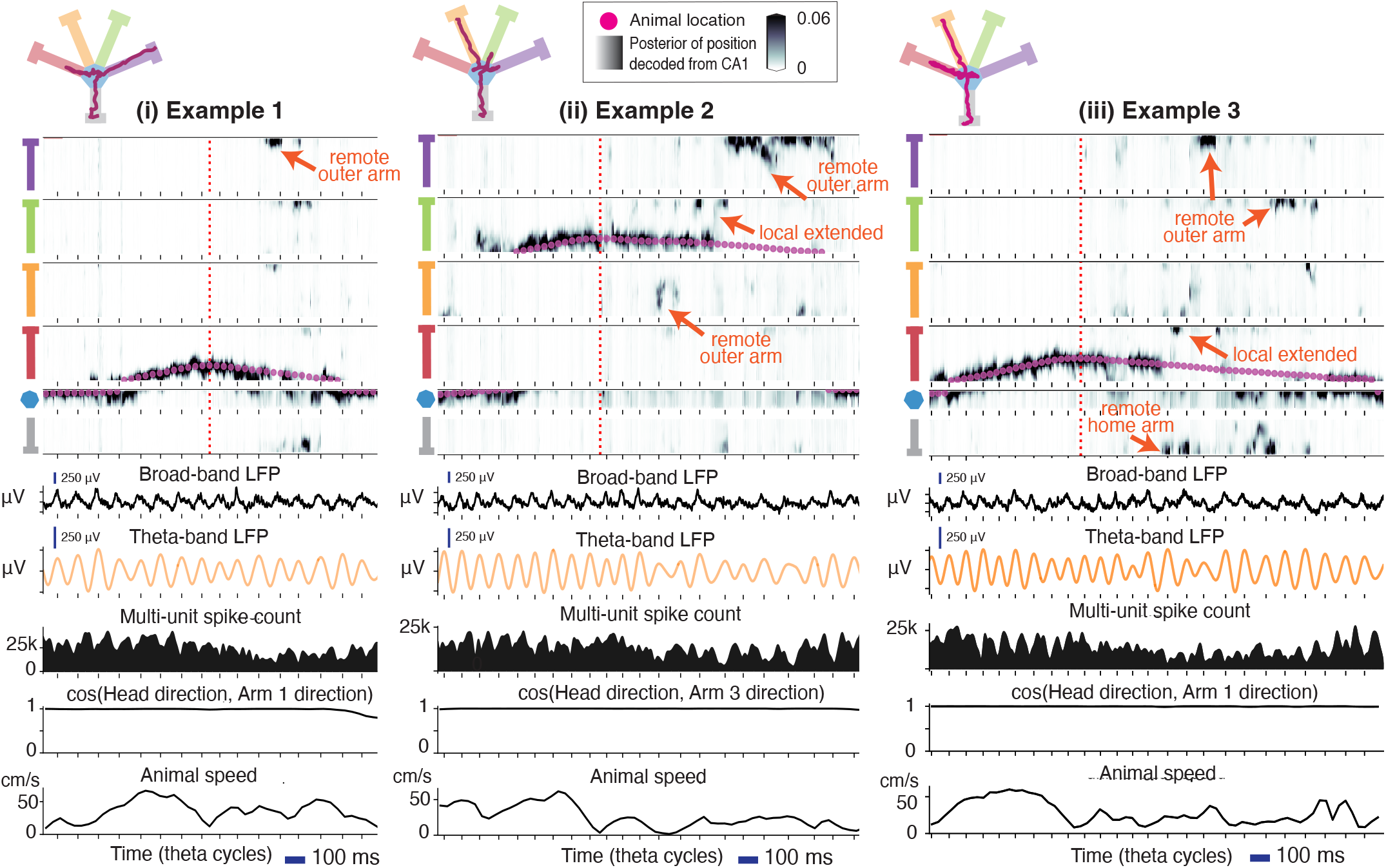
Hippocampal spatial decoding and animal head direction during changes-of-mind. (i)A COM trial from Rat M. From top to bottom: 2D overhead view of maze illustrating animal location, hippocampal spatial decoding, broad-band local field potential (LFP), theta-band LFP, spike count of multiunit activity, cosine of the angle between head direction and the direction of the arm the animal is occupying at the time of COM, and animal head speed. Remote representations by the hippocampus include arm 4, arm 3 and the home arm. Same event as in Figure 1F(i), except only the 2 seconds around COM is plotted. (ii)A COM event from Rat J. Remote representations include the end of the current arm (local extended), arm 2 and arm 4. Same event as in Figure1F(ii). (iii)Another COM event from Rat J. Remote representations include the end of the current arm, the home arm, and arms 3 and 4.

We first focused on representations within the COM arm before stopping. Previous work has shown more extended representations during times of uncertainty (*18*), and plots of the animal position vs. the peak of the posterior distribution from the decoder suggested the possibility of more extended representations before COM events as compared to the same locations in run-through trials (Figure 3A(a), A(b), S3). We quantified this observation by computing, for each COM trial, the maximum distance between the peak of the posterior distribution and the animal’s actual location.

**Figure 3.**
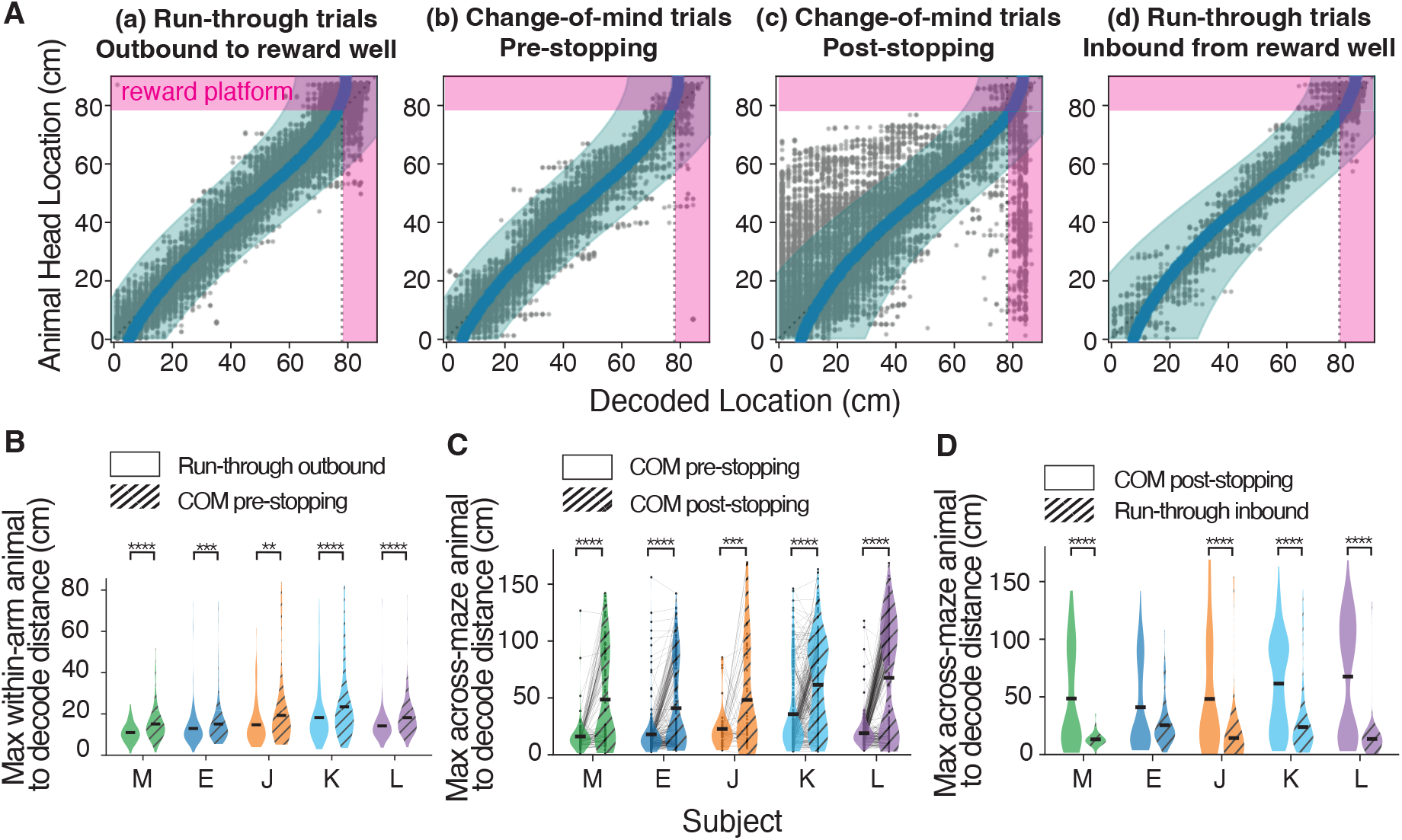
Hippocampal representations of alternatives are expressed after stopping. A. Plotting decoded location from the hippocampus against animal head location, on (a) run-through trials during outbound runs to outer wells preceding the (b) COM trials prior to stopping and (c) after stopping, and on (d) run-through trials during inbound back to home. Shaded teal region denotes confidence interval of decoded location when animal is at various maze arm locations. These intervals are used for local-extended content detection in Figure 4. The confidence intervals in (a) and (d) are obtained by fitting all run-through trials preceding the COM trials on that day during outbound (a) or inbound (d) runs. The confidence intervals in (b), (c) are the same as in (a), (d) respectively. Magenta region highlights the reward zone. Data in these plot come from 1 day of data (day -2) from Rat L. B. Comparison of maximum per-trial within-arm head-to-decode distance for outbound run-through trials (hollow fill) and COM trials in the pre-stopping period (hatched). The regions of maze segments in which the distance is calculated are matched to account for the fact that COM trials do not cover the entire outer arm region. Rank-sum tests, p<0.01 for all rats. C. Comparison of per-trial maximum head-to-decode 2D maze distance between the pre-stopping period and post-stopping period on COM trials. Paired t-tests, p<0.001 for all rats. D. Comparison of per-trial maximum head-to-decode 2D maze distance between the inbound run-through trials and the COM post-stopping period. Regions of maze segments in comparisons are matched to account for the fact that COM trials do not cover the entire outer arm region. Rank-sum tests, p<0.0001 for all rats but E.

We found that there were more extended representations preceding stopping on COM trials than run-through trials (Figure 3B, ranksum test, p < 0.01 for all rats). Further, the longer decoded distances were predictive of whether the trial would have an upcoming COM event (Mixed logistic regression, mean multiplicative effect (95% CI): 1.06 (1.05, 1.08), p < 1e-20). However, the representations did not typically reach the end of the current arm or to other arms. Thus, these representations could contribute to the decision to stop, but it is not clear how they could contribute to the subsequent decision of which arm to choose next.

By contrast, after the animal stopped and began reversing direction, we observed representations that extended both to the end of the current arm (“local extended content”) and representations of locations in other arms (“remote content”) during animal locomotion. Plots of decoded location indicated that represented location could “jump” to distant discrete locations (Figure 2, Figure S4, and Figure S3, see below) rather than follow a continuous path extending along the current path (continuous “theta sequences”) (*16, 28, 29*) (But see (*18, 30*)). These more distant representations were much more prevalent following stopping (Figure 3A (b, c), 3C, paired t-test, p < 0.001 for all rats, Figure S5). We also found that post-COM representations were more likely to represent distant locations than when animals were in the same location without a COM (i.e., when the animal had visited the outer arm well, turned around, and returned to the home arm; Figure 3A(d), 3D).

These jumps in spatial representations were associated with reversals of movement direction but are unlikely to be due to backing up behavior. First, prior studies of backwards walking revealed that the theta sequences remain close to the animal similar to those observed during forward running (*20, 31*). Second, we also observed remote content during normal forward running after the animal had turned around but had yet to exit the arm (Figure S6).

### Hippocampal representations of alternatives predict correct choices

As the COM behavior often corrected initially erroneous choices (Figure 1G), we next asked whether remote representations expressed during COM events could contribute to corrective decisions. We first noted that, on average, COM events engaged an increasing number of representations across all five animals (Figure S11A, mixed linear model mean/(CI)/p value for the effect of days 0.03 / (0.01,0.05) / p < 0.005), consistent with greater engagement as animals recognized the conflict between the task rules and their alternation bias.

We found that the presence of local extended or remote arm was, on average, associated with higher likelihoods of correct subsequent choices across animals (Figure 4A), and that when data from all animals were considered, there was a highly significant effect (Figure 4B, mixed logistic regression; point estimate of fold change, (95% CI), p value: 1.50, (1.18∼1.90), p<0.001, see also Figure S11B for analysis by theta cycle). Moreover, decoded representations on COM trials often involved a combination of the home arm, where the animal would typically start each trial, and either local-extended or remote arm representations (Fig. 4C). As each interval of remote representation typically corresponded to a single location (Figure S5), this suggests that the hippocampus serially represent both the start and end of each journey, as though the animal played out segments of complete trials in greatly accelerated form.

**Figure 4.**
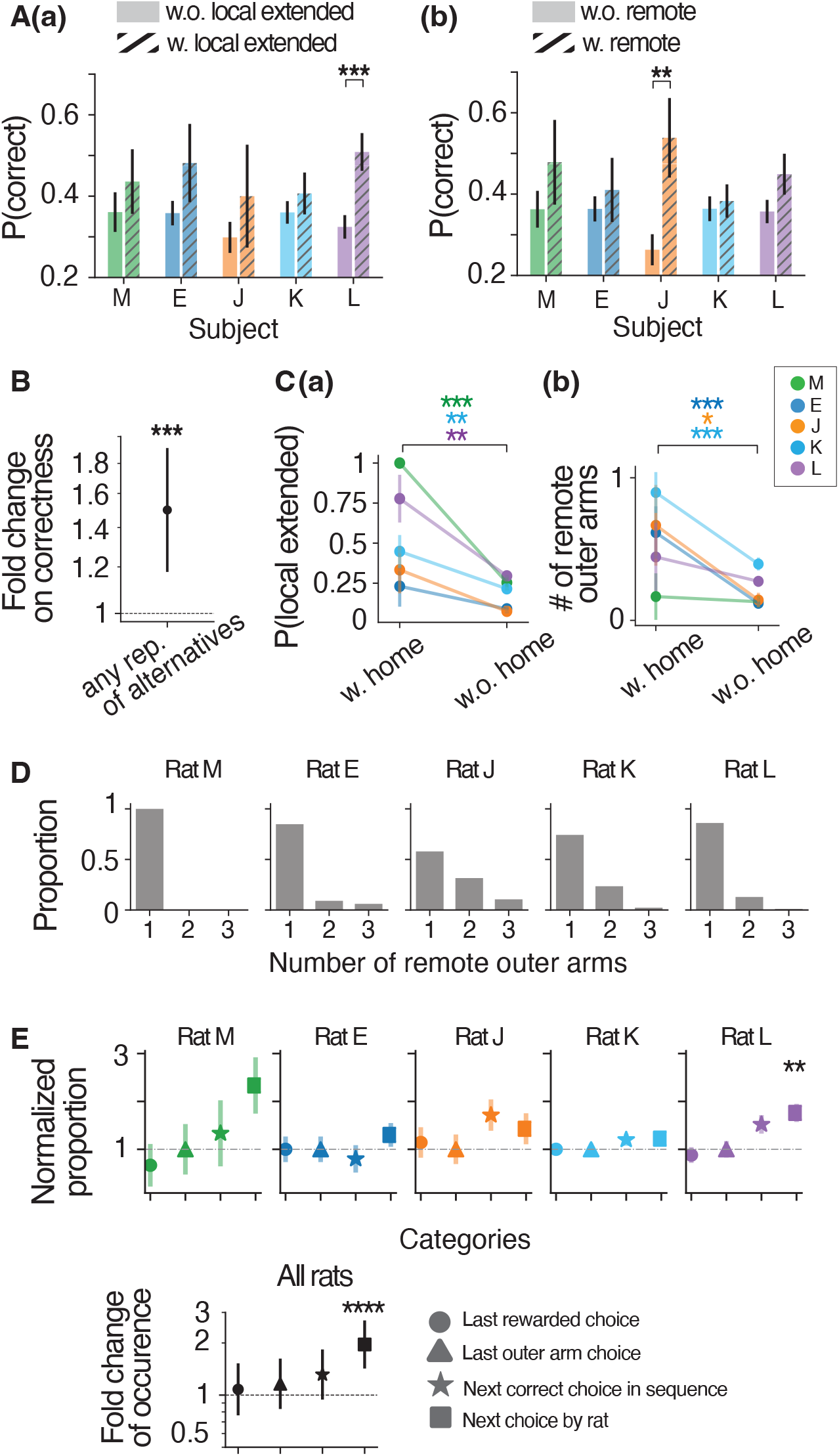
Hippocampal representations of alternatives predict correct choices. A. Probability of the subsequent choice being correct on COM events (a) with no local extended or (b) remote representation. In (a) for “without” n = 113, 242, 133, 250, 280 for rat M, E, J, K, L, and for “with” n = 23, 39, 26, 141, 98. In (b) for “without” n = 97, 254, 144, 300, 262, and for “with” n = 39, 27, 15, 91, 116. Fisher-exact test, p < 0.001 and p <0.01 for rat L in (a) and J (b). Pooling data from all rats with mixed logistics regression: point estimate of fold change on correct rate due to is (a) local extended content is 1.61, 95% CI: (1.22∼2.11), p<0.001, (b) remote content is 1.50, 95% CI: (1.18∼1.90), p<0.001. B.Mixed logistic regression of the subsequent choice correctness onto the existence of representations of alternatives (either remote events or local extended events). Point estimate of fold change on correct rate is 1.50, 95% CI: (1.18∼1.90), p<0.001. C.Probability of observing a local extended event when there are home representations (left) vs none (right). (b) The number of remote outer arms when there are home representations (left) vs none (right). One-sided Fisher-exact tests show COM events with home representation contain (a) larger probabilities of local extended representation: p < 0.01 for rat M, K, and L. and (b) a larger number of remote outer arms in (b), p < 0.05 for rat E, J and K. For both (a) and (b), combining all n = 5 rats, p < 0.032 binomial test. D. Histogram showing the number of remote outer arms that appear in remote representations for each COM event in each rat. E. (Top) Proportion of last rewarded choice (circle), last outer arm choice (triangle), the next correct choice (star), and the next choice by rat (square), in remote outer arm representations during COM events, normalized to last outer arm choice. Error bars denote normalized standard error. (Bottom) Beta coefficients from mixed logistic regression pooling data from all subjects. Fold change due to immediate next choice is 1.96, 95% CI: (1.45∼2.65), p value < 0.00005. n =18, 35, 21, 135, 93 for rat M, E, J, K, L.

We then asked whether the spatial content of these remote representations was related to future decisions. Most often, only one of the three other arms was represented (Figure 4D). If this remote representation serves as a “where should I go next?” signal, then animals would be more likely to visit that arm. To test this hypothesis, for each COM event, we categorized remote representation content into four categories, previous correct arm chosen, previous arm chosen, next arm chosen, or next correct arm chosen.

We found that remote arm representations were predictive of the immediate next choice (Figure 4E, mixed logistic regression for all rats data, fold change (95% CI), and p-value due to rewarded past, immediate past, correct future, and next choice are 1.07 (0.78,1.50), p < 0.7; 1.16 (0.85,1.59), p < 0.35; 1.31 (0.96, 1.80), p < 0.1; 1.96 (1.45, 2.65), p < 0.0001). This remained true with a much less restrictive criterion for identifying remote arm representations (Figure S11C). Thus, COM trials with hippocampal representations of remote alternatives provide predictive information about the upcoming choice well in advance of behavior (the entry into the next chosen choice arm).

### Representations of remote alternatives during COM are concentrated in reward regions

Inspection of individual examples of remote representations during COM events (Figure 2) suggested that, unlike typical theta sequences, COM-associated representations often discontinuously “jumped” to locations at the ends of arms where reward was delivered. This is akin to mental representation of a goal location when the animal is deciding where to go next. To determine whether this was characteristic of a common feature of remote representations during COM events, we quantified the overall distribution of represented locations following stopping.

The results indicated a strong bias toward representations of the ends of the arms in individual epochs and across epochs for all animals (Figure 5A, linear regression of log(proportion), arm end is the largest contributing spatial bin for remote events for all 5 rats; p <0.01 for rats M, E and L, p < 0.05 for rat J, p < 0.1 for rat K, p < 0.0001 for all rats combined in a mixed linear model, mean (95% CI) of effect size is 6.05, (4.23, 7.87)). This jump to the reward area of the arm is also seen in local extended content (Figure S3). Thus, COM events in this task engage hippocampal representations concentrated near reward regions.

**Figure 5.**
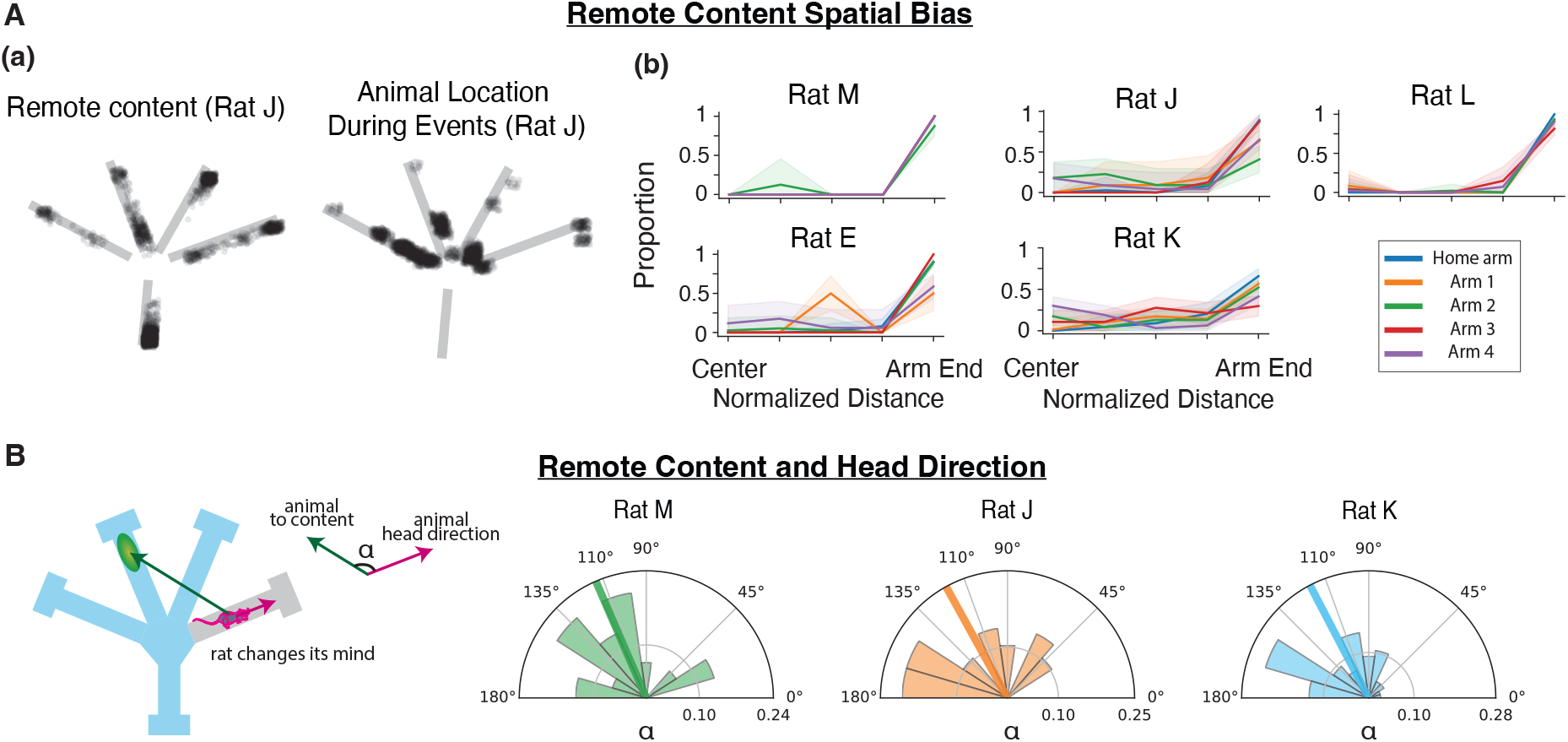
Representations of remote alternatives during COM are concentrated in reward regions located behind subjects. A. 2D scatter plot of all 2 ms time bins of remote event content (left) along with rat location (right), data from rat J. (b) 1D spatial histogram of remote content in each arm across 5 rats. Each arm segment is divided into 5 equally spaced bins. B. Schematic (left) and the histograms (right) of the angle between the animal heading and the direction going from the animal pointing to the content. Bold line shows the mean angle.

Finally, we asked whether the animals tended to orient toward these remote options as they considered them. In three of the animals we had access to a gyroscope readout that, when combined with the overhead video, allowed us to very accurately reconstruct head direction. We therefore measured the angle between the animal’s head and the location of each detected remote representation.

We found that most of the remote representations were located at large angular deviations from the head corresponding to locations behind the animal (Figure 5B). We did not monitor eye position, but as the estimated width of the rat binocular field ranges from ∼40 degrees to ∼110 degrees, depending on head tilt (*32*), a substantial proportion of the remote content is likely to fall outside of the binocular field (proportion of events at angle > 110 degrees: Rat M: 0.67, Rat J: 0.55, Rat K, 0.54).

## Discussion

Change-of-mind behavior is a hallmark of flexible decision-making, where internal processes monitor and reevaluate decisions, weigh alternatives, and select a new choice. Here we developed a paradigm with multiple options in which we simply let the animal adapt to an environment that rewards a foraging strategy that conflicts with pre-existing preferences, with no explicit cues to elicit “second thoughts”. This paradigm prompted many self-initiated COMs correcting for the introduced mismatch in foraging preferences and allowed us to study the internal processes that could support COM decisions.

Strikingly, we found that the underlying neural processes are temporally separable. Using animal stopping as a behavior marker, we found an increase in the extent of nearby spatial representations prior to stopping, consistent with the engagement of these representations under uncertainty (*18*), but very few alternative-choice representations. By contrast, alternative-choice representations emerged robustly after stopping, consistent with a discrete transition from error monitoring to alternative selection.

Beyond this temporal dissociation, we uncovered an unexpectedly rich repertoire of hippocampal spatial representations tied to behavior. These representations were concentrated at task-relevant reward regions and often serially encoded locations associated with the start and end of trials. Moreover, the expression of remote spatial representations was predictive of better choices, and the content of these representations was predictive of the specific choice that would be made next. A recent report of predictive remote representations during immobility theta (*33*) suggests that these remote representations can be expressed across movement and immobility states, and a recent report from our group suggests these representations could also contribute to updating memories related to remote locations (*18*). Taken together, these findings demonstrate a rich capacity for theta-related remote representations that stands in sharp contrast to classical theta sequences observed during locomotion, which primarily reflect the animal’s immediate surroundings rather than distant alternatives (*34–38*).

These results provide new insights into the neural processes that support flexible self-correction. Our results suggest that error monitoring and action selection are neurally dissociable computations, with detection of errors likely occurring outside of the hippocampus while action selection can engage remote hippocampal representations to evaluate alternatives.

## Acknowledgments

Shijie Gu would like to acknowledge the input from Emily L. Mackevicius in early stage of the project, Joni Wallis for suggestions on analysis, Vijay Namboodiri for suggestions on task design, Sam Bray and Chris Broz for assisting with data upload, Katherine Wadhwani for assisting with data collection, Viktor Kharazia for assisting with histology, Vanessa Bender for editing the manuscript, and Matt Rienzo for editing the manuscript.

## Funding

UC Berkeley – UCSF Joint Graduate Program in Bioengineering (SG)

Simons Foundation grant 542981 (SG, CL, AKG, MEC, LF)

K99: NIA K99AG068425 (AKG)

Howard Hughes Medical Institute (RN, ELD, KK, LF)

## Author contributions

Conceptualization: SG, AKG, LMF

Methodology: SG, AKG, CL, RN, ELD, LMF, KK

Data acquisition: SG, CL, MEC

Investigation: SG, LMF, KK

Funding acquisition: LMF

Writing – original draft: SG

Writing – review & editing: SG, AKG, CL, ELD, LMF, KK

## Data, code, and materials availability

Data is available for download from Dandi, the BRAIN Initiative supported data archive, at https://dandiarchive.org/dandiset/001836. Codebase for analysis is archived at: https://zenodo.org/records/20371883. Instructions for downloading data as well as running reproducing analyses are available at: https://github.com/shijiegu/Gu2026_docker.

## Supplementary Materials

Materials and Methods

Figs. S1 to S12

Tables S1

References (*1–51*)

Movies S1, S2, S3: Hippocampal spatial decoding during change-of-mind. Movie S1, S2, and S3 correspond to the example (i), (ii), and (iii) in Figure 2.

Data S1: DANDI:001836: https://dandiarchive.org/dandiset/001836

