## Supplementary material for "Naturalistic behavior and self-generated neural activity predictive of self-correction": Methods and Supplementary figures

**Method**
**Resource Table**

| **Resource** | **Source** | **Identifier** |
|---|---|---|
| **Deposited Data** | | |
| Neural and behavioral dataset | This paper | See below |
| **Experimental models** (Organisms/strains) | | |
| Male Long Evans rats | Charles River | RRID: RGD_2308852 |
| **Hardware and devices** | | |
| Maze design | This paper | Figure S11 |
| Maze welding | Tap Plastics | https://www.tapplastics.com/ |
| 12.7 µm nichrome wire | Kanthal/California Fine Wire | Item: PX000004 |
| Microdrive body | This paper, modified from (*39*) | |
| Data acquisition system: MCU, ECU, headstage | SpikeGadgets | https://spikegadgets.com/spike-products/ |
| **Software and algorithms** | | |
| Database | Spyglass | See below |
| Custom Analysis Code | This paper | See below |
| Hippocampus place decoder | Denovellis et al., 2020 | https://github.com/Eden-Kramer-Lab/replay_trajectory_classification |
| Gyroscope and accelerometer integrator | Denovellis et al., 2025 | https://github.com/edeno/trodestrack |

**Data and code availability** Data is available for download from Dandi, the BRAIN Initiative supported data archive, at https://dandiarchive.org/dandiset/001836. Codebase for analysis is archived at: https://zenodo.org/records/20371883. Instructions for downloading data as well as running reproducing analyses are available at: https://github.com/shijiegu/Gu2026_docker.

**Experimental model and subject details**
Data from five male Long-Evans rats (Charles River aged 4-12 months; 450-550 g) were included in this study. One of these animals had previously been used for recordings in a different environment (*40*). One subject is excluded from analyses due to its poor recording quality. Animals were kept on a 12-hour light-dark cycle (lights on 6am - 6pm) and initially had ad libitum access to food (standard rat chow) in a temperature- and humidity-controlled facility. Beginning one week prior to and extending throughout behavioral training and data acquisition, rats were singly housed and food restricted to a maximum of 80%-85% of free-feeding weight.

All procedures were approved by the Institutional Animal Care and Use Committee at the University of California, San Francisco.

**Behavioral training**

The behavioral room included a sleep box (12”x12”x16” walls) and a maze environment equipped with automated ports in a room with decorations on all four walls. All behavior experiments are conducted in dim light. Task control was automated by an Environmental Control Unit (SpikeGadgets) running custom behavioral code written in Python and Statescript (SpikeGadgets). Reward milk wells are made of an internal white LED, an infrared photogate and a reward delivery tube connected to a syringe pump (Braintree Scientific) that can dispense precise quantities of reward (Nestle Carnation Evaporated Milk + 5% sucrose) when an IR beam break is detected (*41*).

The maze takes about 1.5 m x 1.5 m of space, consisting of a home arm (0.4 m), leading to a small central platform (maximal radius 0.2 m) that is equidistantly extended into four choice arms (96 cm each) (Figure S1A). The whole maze is enclosed by 40-cm-tall walls joined to the maze at a 75 degree angle (in each arm, the width is 8 cm at the base, enclosed by slanted walls extending to 22cm wide); the regions of which that are over arms and reward stations are transparent; the ground piece of the maze is black. From the maze, subjects could see room cues, including wall markings, the door, the sleep box, and a desk. Each arm ends in a reward station (18 cm x 22 cm). The distance from the beginning of the arm to the end of the reward station is 96 cm. The maze was designed in CAD. The maze is then custom-built by TAP Plastics, San Francisco (see Resource table). All pieces are saw cut and router cut from acrylic, and are of thickness 1/4-3/16”; maze walls are plastic welded to the ground pieces.

Rewards are delivered based on a hidden spatial-temporal structure among the 5 reward wells in the maze (home well, outer choice wells 1-4). The home well dispensed 100 µL of milk while the outer ports dispensed 400 µL following a correct nose poke. Subjects perform the complete task as follows: each self-paced trial is initiated by a nose trigger at the home well and receipt of a reward. After reward consumption at the home well, all outer choice ports are illuminated, and the rat visits a single arm before returning to the home port to initiate the next trial. At any given time, only one arm port (“goal arm”) would provide a reward; the others would provide nothing. An arm becomes a goal arm after the reward at the specific previous goal arm is obtained. For example, suppose a correct sequence is: home – 1 – home – 4 – home – 2 – home – 3 – home – 1. The reward at arm 4 is dispensed only after the animal has already obtained a reward at arm 1 and has poked the home well before poking the well at arm 4. During learning, rats usually sample multiple nonrewarded arms before finding the goal location. If the animal visits an incorrect outer arm well, for example, choosing arm 2 after receiving a reward at arm 1 and then the home arm, the next correct arm remains fixed as arm 4. There are no cues that indicate the next correct arm.

In our pilot behavior cohort we found that naïve rats show a tendency to alternate between the left and right side of the maze (switch-side transitions). This was also observed in our experimental cohort. To encourage conflict between this tendency and the task rules, we chose 4 arm cycles with an equal number of same-side and switch-side transitions. There are 6 possible 4-arm cycles, which can be grouped into 3 sets omitting the home visits in between: (A) 1-2-3-4-1 and 4-3-2-1-4, (B) 1-2-4-3-1 and 3-4-2-1-3, and (C)1-3-2-4-1 and 4-2-3-1-4. Set A contains 1 side-switching transition (4-1 or 1-4); set B contains 2 side-switching transitions (2-4/4-2 and 3-1/1-3); set C contains exclusively side-switching transitions. Therefore, to achieve a balanced number of transitions of same-side and switch-side transitions, we only tested 4-arm cycles in set B in our rats.

Before running subjects on this task, they are trained to obtain milk through poking reward wells and to initiate a trial at home and finish it at the outer choice well. The first two stages of this training occur before the neural implant (see below). Stage 1 is the exposure of the condensed milk to rats. Milk droplets are randomly scattered in their home cage to familiarize rats with milk. This stage lasts about 2 days with milk droplets added daily. After Stage 1, animals are food-restricted to 85% of their body weight for Stage 2. In Stage 2, we train the subjects to alternate between two reward wells in a linear track environment. The linear track is set in the same room as our main task maze and is oriented along the home-outer well axis. The track is 50 cm long with 60 cm tall opaque walls. We run approximately 3 training sessions of 15 min each per day, with 30-60 min of rest in their home cage in between. Training continued until each animal received more than 50 rewards in 15 min in two consecutive sessions (~5 days). Rats are then given at least 7 days of rest before potential future stages.

In each cohort of 3 rats, the top-performing rat in Stage 2 is selected to be implanted with electrodes. After 10 days of post-op recovery with *ad libitum* access to food, subjects are food-restricted to a maximum of 85% of their free-feeding weight. Subjects are next acclimated to the maze environment and learned to alternate between home well and any of the outer wells. In this stage, we run sessions of 80 trials each with no time limit as long as the rat does not stop for an extended time (10 minutes) on the track. In that case the rat is taken out of the maze and put in the sleep box for 20-40 minutes. We gradually increase the number of daily sessions, from 2 sessions a day to 4 sessions a day. For each trial, the rat needs to alternate between the home well and any of the outer wells. To shape this behavior, a "timeout" white noise sound was triggered if the subject poked any two outer wells without poking the home well in between.

In some cases a rat would alternate between 1 specific outer arm and the home arm. In those case that arm would be blocked for one session to encourage exploration. All rats were advanced to the full task once they have reached the criteria of consecutive 2 sessions of greater than 64 (80%) rewards in a session of a total 80 trials and less than 8 (10%) of timeouts.

After behavior shaping, we run the full maze task. During the first few days of the full maze task, due to the sudden drop in the reward rate from 80% to a low number, Rat E, K, J, and L all became unmotivated so that only 2-3 sessions could be run, and a session would sometimes have at most 12 trials. But as the animals acclimated to the maze environment and started to explore more, the number of sessions was increased to 4-6. Each session was flanked by 30-60 min sleep sessions in the rest box. This acclimation phase took about 3-7 days for each rat. Training continued until behavior plateaued, except in rat M, where only 6 days of data could be collected. Behavior is defined plateaus when the reward rate does not progress in 3 days after the correct sequence becomes the dominant sequence in behavior. We focused on data from 5-11 consecutive days ending on day of the peak performance to span from low to high performance on the target sequence (Figure S1E).

**Neural implant**

Each implant housed 32 (for Rat E) or 64 (for Rat M, J, K, and L) independently moveable nichrome tetrodes (12.7 µm, Kanthal/California Fine Wire), cut at a sharp angle and gold plated to a final impedance of ~200-350 kOhms. The implant was stereotactically positioned such that each of the bilateral cannulas containing 16~32 tetrodes (1.5 mm diameter) was centered at 4 mm AP, ± 2.6 mm ML relative to the skull bregma. A screw placed over the cerebellum served as a global reference during data collection. Subjects were given 10 days of recovery after surgery, with ad libitum food and daily tetrode adjustment until the dorsal CA1 cell layer was reached. We note that through post-experimental brain slice histology, the implant of rat K was discovered to be placed slightly anterior in coordinates and some electrodes was putatively placed in CA3 instead of CA1. In all rats, we included all electrodes with visible SWRs. Thus, in the case of rat K, these putative CA3 electrodes were included as well. Cell yield remains high throughout recording (Figure S12).

**Data collection and processing**

Continuous 30 kHz neural data, environmental events (port beam breaks, lights, reward delivery), online position tracking of a head-mounted LED array, and 23 Hz video were collected using the headstage/MCU/ECU system (SpikeGadgets) and two PTP enabled Allied Vision cameras, one over the maze and one over a sleep box. Data were converted to NWB format and stored in the relational database in Spyglass (*42*). All analyses were performed using common code in Spyglass and additional custom code.

**Electrophysiology**

Two electrodes in each implant are left in the corpus callosum.

Local Field Potential (LFP) was extracted from the first functioning channel from each tetrode, by filtering the reference-subtracted continuous signal between 0 - 400 Hz (0-400Hz pass, 400-425Hz stop). SWR hippocampal LFPs are obtained by first subtracting from them the raw continuous signal of one of the channels of the ipsilateral corpus callosum electrode, followed by band-pass filters to produce SWR LFP (150-250Hz pass, 140-260Hz stop). We use filters implemented by the ghostipy package(*43*). Spiking events are obtained by first subtracting from them the raw continuous signal of one of the channels of the ipsilateral corpus callosum electrode, followed by band-pass filtering 600-6000 Hz, using spikeinterface (*44*) filters. Spike events were detected when the voltage exceeded 100 μV on any channel of a tetrode.

**Position tracking**

Rats wear head-mounted LED array pair (front and back arrays of 5 LEDs each) and their locations are tracked by Trodes software (Resource table) through 28 Hz video collected alongside electrophysiology data. These raw LED position data were further processed in the following steps: (1) remove unlikely jumps by replacing these data points with NaN: when the instantaneous speed of a LED is above 300 cm/s or when 2 LED rings are greater than 9 cm apart; (2) linear interpolation of NaN (3) smoothing by applying Gaussian smoothing with a kernel size of 0.125 s. Each LED array position trace was then linearly up-sampled to 500 Hz for place decoding (see below). Animal position was then calculated from the centroid of the front and back LEDs. This smoothed and upsampled position trace is used to calculate animal head velocity.

**Position linearization**

The 2D position data processed as detailed above are projected to 1D data along the arms. For the center platform position data, they are projected to the midline going from the base of the home well to the tip of the center platform (*45*).

**Head direction tracking**

Due to the reflection of the LED on the wall of the maze, the 2D tracking of the animal location, as well as head direction, is not accurate. We note that the 1D tracking was accurate as we collapse the reflections on the maze walls onto the dimension along track length.

We utilize an extended Kalman filter state space model to combine data from 3 noisy sensors: the overhead position tracking, the gyroscope, and the accelerometer on the 256-channel headstages that Rat M, J, K, and L have on their implant. The model predicts position and derivatives based on motion dynamics derived from the inertial sensors, which are sampled at a higher rate than

the camera (104 Hz vs. 23 Hz). These predictions are updated using camera-based observations. By weighting each source according to its uncertainty (via the Kalman gain), the model provides robust position estimates during brief periods of unreliable visual tracking. We set measurement noise for inertial sensors based on device characteristics and tuned observation noise for camera-derived positions empirically. The tracking package is available at:
https://github.com/edeno/trodestrack

### Theta LFP

We used LFP recorded from corpus callosum. We filtered the LFP with a bandpass filter (5-11Hz pass, 4.5-12Hz stop) to produce theta LFP, which was then Hilbert transformed using a 5-11 Hz band pass filter with 4.5Hz and 12 Hz stop to obtain phase. In all analyses pertaining to theta except Figure S9 and S11, theta LFPs from corpus callosum were used. For Figure S9 and S11, we used multi-unit activity (MUA) derived theta for all rats due to the electrodes in corpus callosum in Rat K and L have slightly sunk into the CA1 cell layer. To derive theta phase from MUAs, we followed the procedure as in (*46*). MUAs are smoothed with a Gaussian kernel of size 40ms. Peaks of MUAs were detected with the scipy function find_peaks() with a minimum distance of 100 ms. Phase in between peaks were linear interpolated.

### Quantification and statistical analysis

#### *Behavioral analysis*

Trial start and end time marked by the subject poking at home well and outer choice wells, along with reward deliveries, were extracted from behavior log files (Statescript; SpikeGadgets).

#### *Change-of-mind (COM) trial detection*

COM trials were found by detecting trials in which the animal traversed multiple outer choice arms. A threshold $\theta$ was used to select trials in which the animal's head traversed more than $\theta$ proportion of any other arms than the final arm the animal chose. In Figure S1, we show $\theta$ = 5cm, 10cm and 20cm. In the rest of the paper, we use $\theta$ = 10cm for all analyses.

#### *Change of mind time detection*

We *define* the COM time to be the moment the rat stops advancing forward in a choice arm. We operated on smoothed and upsampled 1D position tracking data. For each time bin animal spent in outer arms, additional smoothing was applied by the binary_closing function from Python Scipy ndimage package (*47*). Consecutive intervals of moving in the negative direction longer than 100ms were considered as reversing. The consecutive window size was empirically set to be a short period of 100ms to allow the identification of vacillation trials in which the animal moves back and forth multiple times. These vacillation trials do occur in multiple subjects but are rare. Each arm choice in which the animal reversed course was considered a COM event, and a trial that contained at least one COM event and animal trajectory into one COM arm for more than 10 cm is considered a COM trial.

*Identification of nearby trials that are not change-of-mind trials*

To generate control data in which the animal did not change its mind, we use trials near COM trials. Given a COM trial, we randomly select trials that were within +/-3 trials from the change of mind trials. In certain analyses, we also controlled for reward — in this case, the randomly selected trial was also a rewarded trial. In some cases, it was not possible to find eligible nearby trials in some sessions. For example, when the animal had learned the task, it was sometimes impossible to find a nearby trial that was not rewarded, therefore the control dataset and the COM dataset do not always have the same number of trials, but these cases are rare.

*Spatial decoding*

We use a state space model (*45*), without spike sorting, to simultaneously decode the "mental" spatial position of the animal and determine whether the position was consistent with a spatially continuous or fragmented movement model. The model relies on an encoding model that builds a conditional probability density estimate of spike amplitudes given positions and decodes by inverting the conditional probability density. The difference between the state space model and the more traditional memoryless Bayesian decoder (*48*) is the inclusion of the latent variable that models the type of spatial trajectory, allowing us to identify a larger set of coherent spatial content not restricted by linear regression. Figure S2 includes decoding example snippets from all rats with the state space decoder and the memoryless Bayesian decoder side-by-side.

We decoded each behavior session independently. For the encoding model, we used all spikes recorded during that session when the rat was moving > 4 cm/s, including those from both pyramidal cells and putative interneurons. We decoded spiking activity for all times during the behavioral session.

To build the encoding model, linearized positions were first binned into 2 cm bins within each maze segment. For each multiunit spike (amplitude > 100 μV on at least one tetrode channel), we associate its waveform features (the amplitudes on each of the 4 tetrode channels at the time of the maximum detected spike amplitude) with the linearized location at the time of the spike using kernel density estimation. We filled the amplitudes on dead channels, if any, on tetrodes, with zeros. Denoting time bins by index $k, k \in \{0,1,2, \ldots T-1\}$, where $T$ was the total number of time bins in this session, linearized position at time bin $k$ by $x_k$, spike features, also called marks of all $N_{k,j}$ spikes at time bin $k$ from electrode $j \in \{1,2,...J\}$ , where $J$ is the total number of tetrodes in CA1 cell layer, by $\{\vec{m}_{k,j}\}^{N_{k,j}}$, $\vec{m}_{k,j} \in \mathbb{R}^4$ (since a tetrode has 4 channels), we obtain the following joint probability for time bin $k$:

$$P(\{\vec{m}_{k,j}\}^{N_{k,j}}, x_k) = \prod_{n \in N_{k,j}} P(\vec{m}_{k,j,n}, x_k)$$

where all spikes contribute independently.

This quantity of the model is fit from all run data (2D headspeed > 4 cm/s). The kernel density estimators used a 24 μV bandwidth Gaussian kernel for spike waveform features and 3 cm kernel for position.

In addition, denoting the movement type at time by

$$I_k \in \{\text{continuous, fragmented}\}$$

we can further define a movement model $P(x_k, I_k \mid x_{k-1}, I_{k-1})$. For example, $P(X_k = b \mid X_{k-1} = x_{k-1}, I_{k-1} = \text{fragmented}) = \frac{1}{B}$, for all spatial bins $b$ (where $B$ is the number of spatial bins) conveys the idea that in the fragmented state, the spatial location can jump to any location. In this model, the movement model, marginalizing out the space variable produces the movement variable switching dynamics $P(I_k, I_{k-1})$, which quantifies how often a stationary state transitions to a continuous state etc. The movement model details are referred to ref (*45*). This quantity of the model is hard-coded.

This movement model, in conjunction with the encoding model expressed as the joint probability of mark and spatial location allows us to obtain the conditional probability of spatial location and movement type given mark, which means we can decode place and classify movement type at the same time. Starting with $P(X_0, I_0 \mid \{\vec{M}_{0,j}\}^{N_{0,j}}) \propto P(\{\vec{m}_{0,j}\}^{N_{0,j}}, x_0)$, which means a uniform prior on movement types, we apply the forward algorithm for $k \in \{1,2, ... T-1\}$ to obtain causal position decoding:

$$P(X_k, I_k \mid \{\vec{M}_{k,j}\}^{N_{k,j}}) \propto P(\{\vec{m}_{k,j}\}^{N_{k,j}}, x_k) \sum_{I_{k-1}} \sum_{X_{k-1}} P(x_k, I_k \mid x_{k-1}, I_{k-1}) P(X_{k-1}, I_{k-1} \mid \{\vec{M}_{k-1,j}\}^{N_{k-1,j}})$$

We decode using a time window of 2ms bins. All decoding analyses are conducted on the obtain causal position decoding. For a memoryless Bayesian decoder, we simply replace the second term on the right-hand side with 1 to ensure every time bin is estimated independently. Due to the lack of information from nearby windows, we use a time window of 20 ms and an overlapping window of 10ms and decode all windows independently.

*Change of mind triggered decoded location verse head direction (Figure 3)*

In Figure 3A, we aimed to characterize the decoded position within the arm and thus only the decode in the arm segment where the animal changes its mind is shown. Moments where the sum of the decode posterior in the change-of-mind arm was less than 0.2 are not plotted. To compare run-through trials with COM events, for each COM we randomly selected a nearby run-through trial for each COM. Because the decode-animal location deviation is a function of how much the animal has advanced into the outer choice arm, to match distribution, in calculating the deviation from run-through trials, a region is randomly selected from the pool of COM events and only deviation from the region was calculated. Because COM events versus a randomly selected trial is not necessarily a paired trial-by-trial comparison, we use ranksum tests to compare the two groups, while for pre-post COM events, we use a paired comparison – paired t-test.

*Identification of trials with large decode to body position distance during theta.*

We label contents that extends to the end of the current arm the animal was occupying at the time of the COM as "local extended content" and content that jumps to other arms as "remote content".

To detect local extended content, a canonical place coding model – decoded location as a function of how far into each outer arm the animal is – is fit by one Gaussian Process (GP) for mean estimation and another GP for deviation estimation. For each day, for all *m* COM trials, we randomly select roughly a total of *m* non-COM run-through trials as control trials. For each of the *m* control trials, we obtain a list of paired tuples of *{x, y}_i* when the animal is moving with 2D head speed greater than 4cm/s, where *x* is the animal body location, *y* is the location decoding from hippocampus, with i indexing timestamps, which are processed to be sampled at 500Hz (see above, *position tracking and position linearization*). All tuples from *m* trials are collected and 1000 of the data points are subsampled (for computational efficiency) to fit the two GPs. The fitting process is done with Python sklearn package GaussianProcessRegressor() and associated functions (*49*). For both GPs, we use an RBF kernel with kernel size of 5 cm, and alpha is set to the square of the standard deviation of the decode to animal distance across each day's data. We obtain the confidence interval as the mean trace plus/minus 6 times average deviation. The teal shading region plotted in Figure 3A are the mean plus/minus 6 times average deviation. Continuous time intervals with longer than 30 milliseconds (ms) maximum a posteriori (MAP) extended outside of the confidence bound of GP are considered local extended events.

*Quantification of posterior concentration among arm segments*

To quantify if the remote place activity was concentrated in one arm or dispersed in a few arms, we carried out the following analysis. We first focus on 2ms time bins or theta cycles in which

- The animal head speed is greater than or equal to 4 cm/s.
- The animal is in outer arms.
- The sum of the decode causal posterior in the arm the animal is occupied is less than 0.5 to identify moments of nonlocal place representation.

Then for each of the time bins, we count the number of arms with the sum of the decode posterior greater than 0.2 or 0.3. Each count goes into a list and the histogram of the list is shown in Figure S5.

*Identification of remote theta events*

To detect remote content, we first detected continuous time intervals of > 20ms length where the MAP was located in choice arms other than the current arm. The use of MAP is informed by the analysis that the content is not ambiguous – one arm that dominates the decoding for each spike decoding time bin (2ms) (Figure S5A) and for each theta cycle (Figure S5B). We consider only times the animal was moving more than 4cm/s. Each interval further needed to satisfy the following conditions:

- The continuous/fragment decoding had posterior of the continuous state > 0.5 for more than a contiguous interval of 20 ms.
- The position decoding (max or mean of posterior) only goes to one arm segment throughout (so that we can unambiguously parse the position decode), including the current arm (so that we can unambiguously call it "remote").
- The sum of the posterior of the position decoding in the remote arm (identified through max/mean posterior summary statistics) was greater than 0.2.

These parameters are partially referenced from (*50*). In Figure S11(C), we conducted a version of analyses without any of the parameters listed above and found that the results were robust.

*Content / Theta Phase relationship*

For each remote event, we analyzed the time window from 60ms (about half a theta cycle) before the event onset to 60ms after the event onset. Further we restricted analysis to the time the animal was in the choice arms. For each 4ms bin, we obtained 1) the theta phase and 2) the sum of posterior probability of decode outside of the outer arm in which the animal was located. For local extended events, all procedures were the same except that each 4ms bin we obtained the decode to animal distance rather than the sum of posterior.

*SWR detection*

SWRs were detected using a combination of the consensus method (*51*) based on the envelope of combined the ripple filtered (150-250 Hz) trace across electrodes as well as the detection method in individual electrodes (*52*). Each tetrode in the hippocampus CA1 cell layer was first annotated a channel to be used for SWR detection. To obtain the envelope of consensus ripple trace, filtered ripple power envelope trace from the annotated channel from each electrode was squared then summed across all electrodes, followed by Gaussian smoothing of 4ms kernel. Events were detected as deviations in the consensus trace exceeding 2 SD above total session baseline (mean) for at least 15 ms, and exceeding 3 SD in individual tetrode ripple traces for more than 1 channels. Session baselines for the consensus trace or individual traces are all calculated when the animal is moving greater than 4 cm/s and SWRs were only detected during immobility (velocity < 4 cm/s).

*Multiunit activity (MUA) event detection*

All spikes > 100 μV on at least one channel of a tetrode in CA1 cell layer were included.

*Generalized linear models to measure the effect of arm category on remote event likelihood*

Logistic GLMs were constructed using the Logit() and fit() function in the Python statsmodel package (*53*). All COM events with remote contents were included. We note that each COM trial could contain more than one COM event, and here we parsed and fit event by event. For each COM event, any arm that the rat had entered for more than 5 cm before and after was considered the immediate past and next choice respectively. One binary predictor was constructed per behavioral category. For instance, for the future arm predictor: each trial contributed 4 entries (one per arm), with zeros denoting arms that were not the future arm for that trial and one denoting the arm that was the future arm on that trial. The 4 response entries for that trial are whether there exist remote events with spatial content in each of the choice arms 1-4. To pool across animals, one binary predictor was constructed per animal. For example, given we had 5 animals, M, E, J, K, L in the dataset, rat M's predictor is [1,0,0,0,0] and rat L's predictor is [0,0,0,0,1]. The model used the canonical link function. The coefficients associated with each animal category represents the remote representation likelihood baseline for each animal. We used the mean estimate and 95% confidence intervals for the resultant coefficients from the fit function of the statsmodel package. We also confirmed that using robust fit with regularization did not qualitatively change the results. The mean estimate and confidence intervals were then converted them into fold change by exponentiation. Results pertaining to the intercept and animal categories are omitted.

*Generalized linear models to measure the effect of representations of alternatives on correctness*

Logistic GLMs were constructed using the Logit() and fit() function in the Python statsmodel package (*53*). All change-of-mind trials were included. We note that each COM trial could contain more than one COM event, and here we parsed and fit event by event. For each COM event, any arm that the rat has entered for more than 5 cm after the current COM event was considered the next choice and the correctness is with respect to that choice. In Figure 4B, the predictor for each trial is the existence of either local extended content or remote content. The response variable was whether the immediate next choice of the animal is rewarded. To pool across animals, one binary predictor was constructed per animal. For example, given we have 5 animals, M, E, J, K, L in the dataset, rat M's predictor is [1,0,0,0,0] and rat L's predictor is [0,0,0,0,1]. The model used the canonical logit link function. The coefficients associated with each animal category represents the remote representation likelihood baseline for each animal. We used the mean estimate and 95% confidence intervals for the resultant coefficients from the fit function of the statsmodel package. We confirmed that using robust fit with regularization does not qualitatively change the results. The mean estimate and confidence intervals were converted into fold change by exponentiation. The same analysis can be conducted using trials with only 1 COM event. The result does not qualitatively differ.

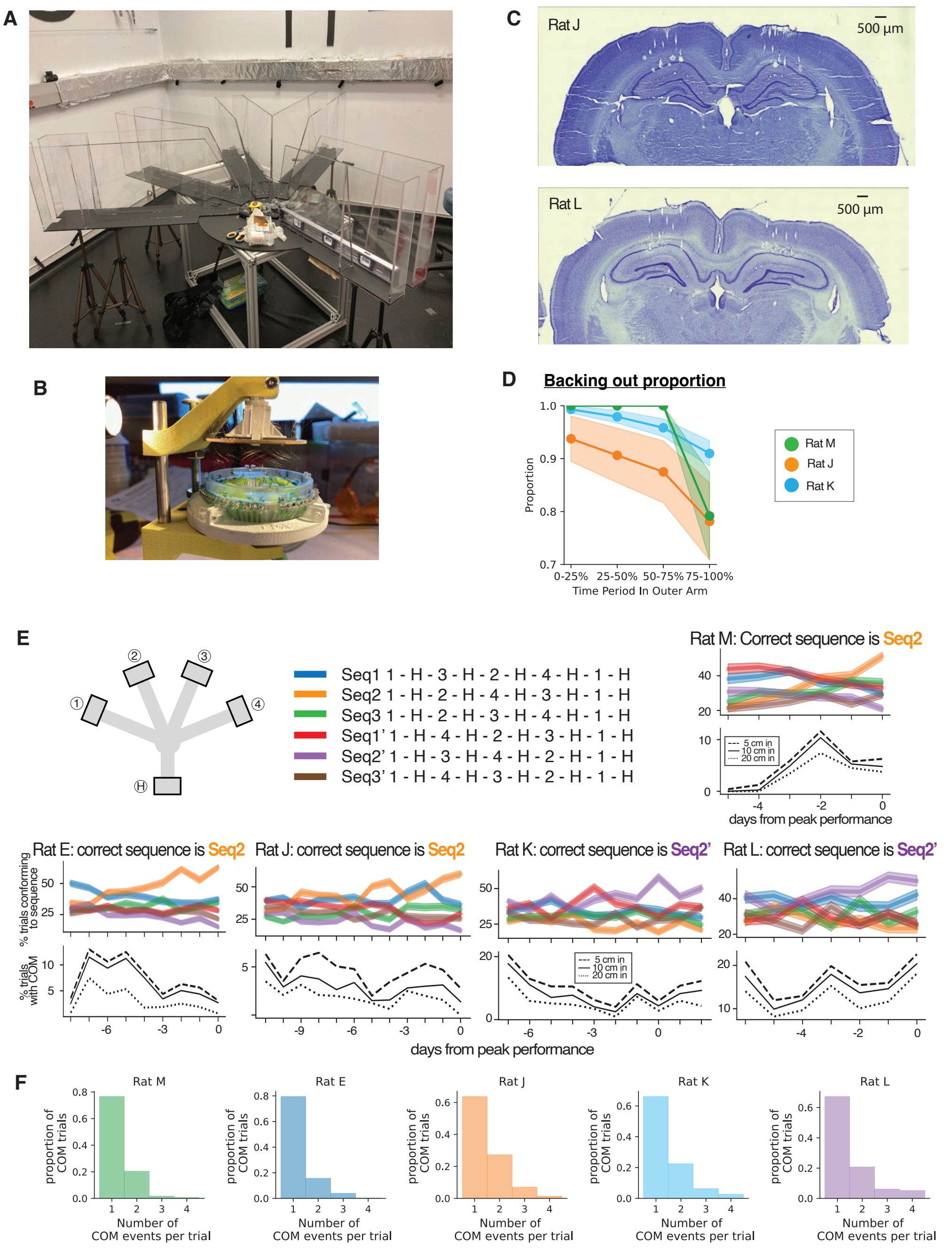


**Figure S1. Additional details of the setup, recording method and behavior analyses.**

A. Image of maze enclosure and the rig room setup. The ruler, bolts, and the scissor in the picture were not part of the maze. The opening at the central platform connecting the home arm to outer choice arms was also enclosed with opaque plastic boards.
B. A 64-tetrode microdrive being assembled.
C. Nissl staining histology from Rat J and Rat L. The tetrode implant locations are visible through the tracks.
D. Quantification of direction of motion following stopping on COM trials. Each trial is split into 4 equally long temporal bins. For each bin we calculated and plotted the proportion of trials facing the reward well.
E. Sequence occurrence in behavior in all rats. In each rat, the top panel shows the percentage of trials conforming to each of the 6 possible sequences (see color legend). The correct sequence for each rat is labeled. The bottom panel shows the percentage trials with COM over the days.
F. Histogram of proportion of COM trials with 1, 2, 3, or 4 COM events. Anecdotally, rat E, the oldest rat in the cohort did the least number of 2 COMs/per trial while rat J, the youngest in the cohort did the most, consistent with the cost of COM.

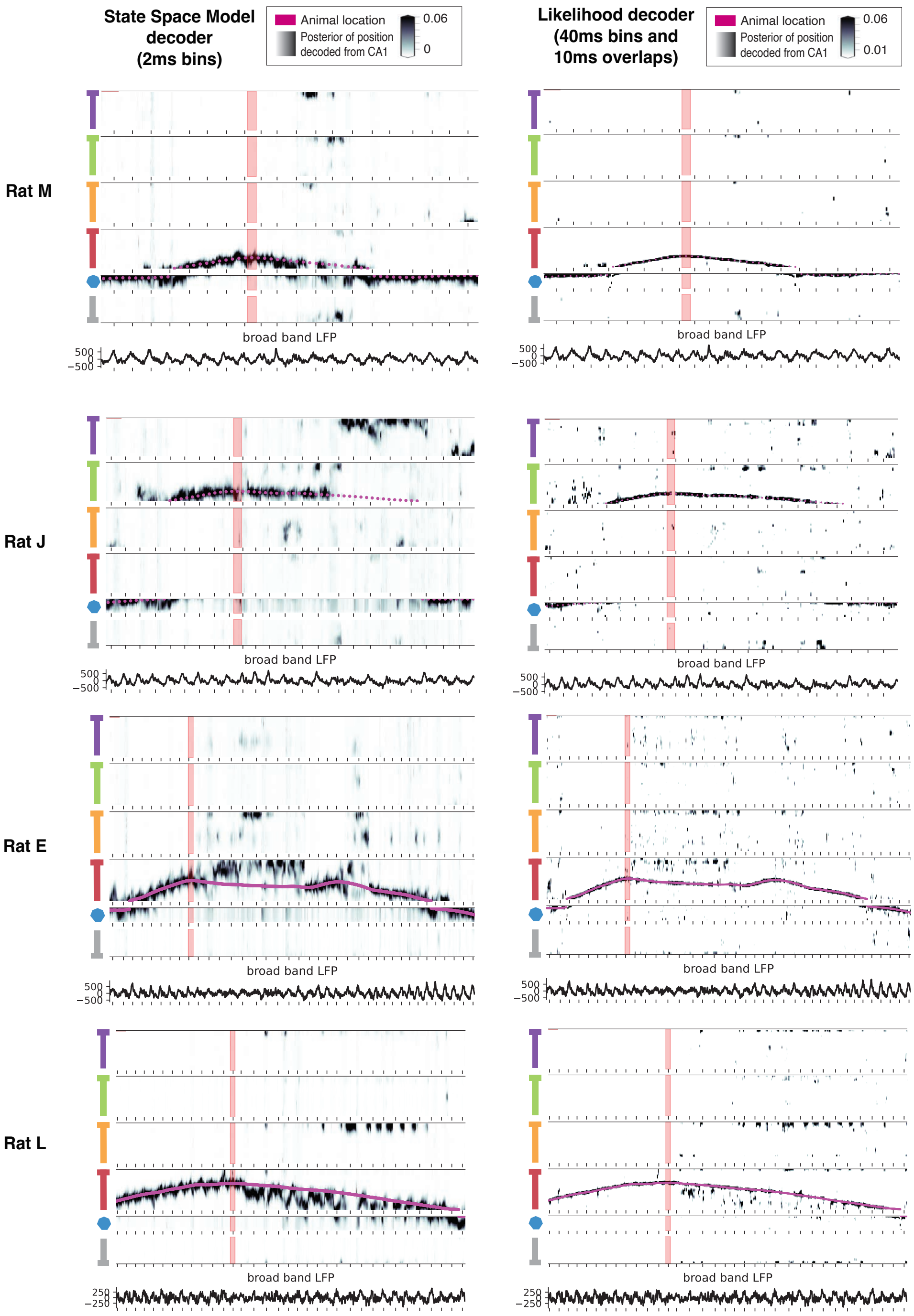


**Figure S2. Addition examples of remote representations around change-of-mind.**

Left panels are states-space Bayesian decoder as used in the main text and right panels show the same examples with memoryless Bayesian decoder. The example from Rat M and Rat J are the same examples in the main text Figure 2, examples i and example ii. For the memoryless Bayesian decoder, we used 20 ms window size and 10 ms window overlap.

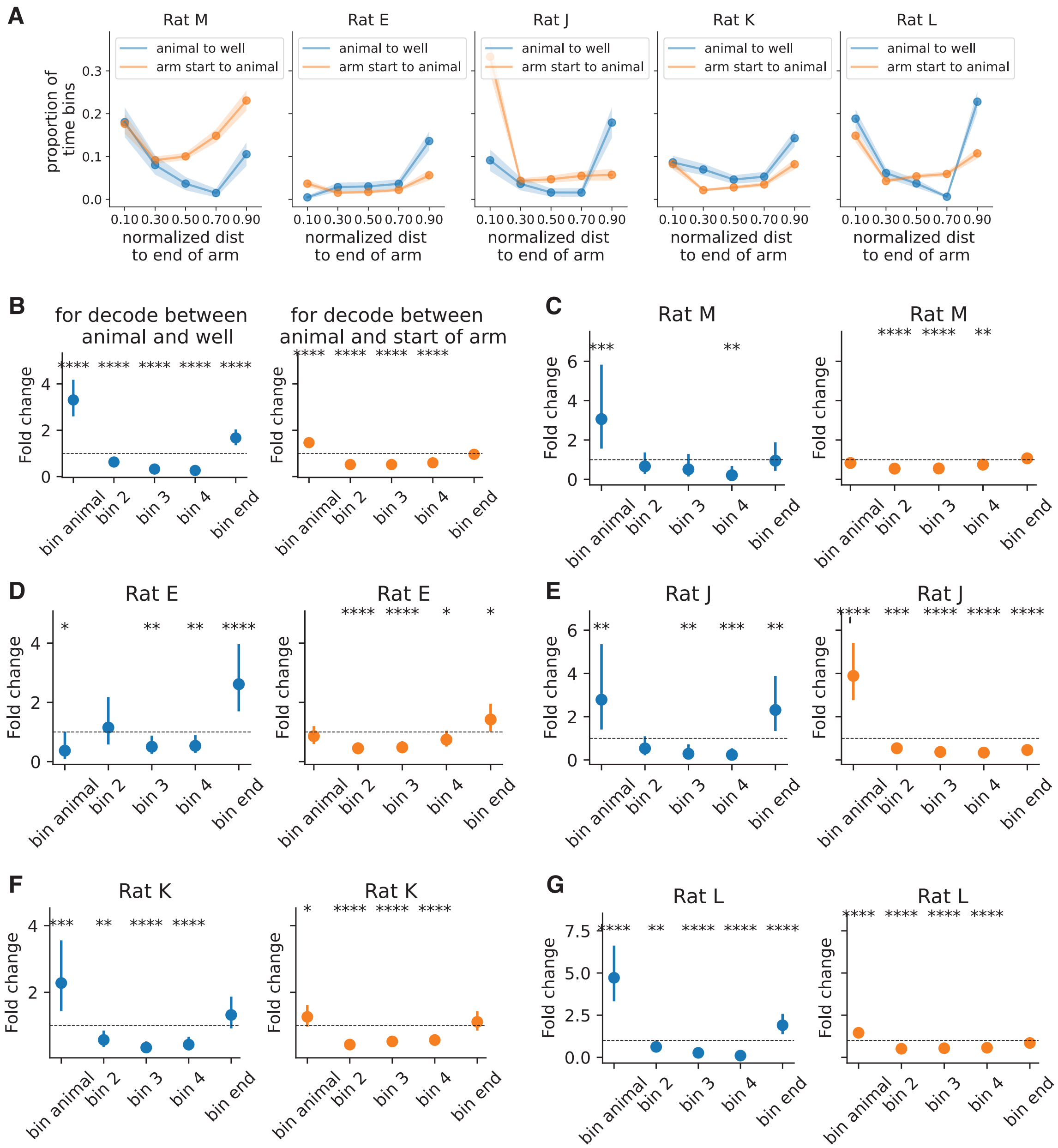


**Figure S3. Proportion of time bins in which the decoded position is close to the animal or to the ends of the arm.**

In Figure 5, we quantified the location of remote representations. In this figure, we quantify the locations of local extended content. We do so by breaking each arm into 10 different bins, 5 in front of the animal (blue) and 5 behind the animal (orange). We normalize the distances so that "bin animal" corresponds to the bin closest to the animal and the "bin end" corresponds to the reward well end of the arm in blue panels and the beginning of the arm in orange panels. (A) Histogram of all decode beyond the confidence interval (fit as in Figure 3). Max posterior in the COM arm when the sum of posterior > 0.2 are considered. Shadings show standard error. (B) Mixed linear effect fit to log(proportion). The reward well bin is over-represented (fold change mean/(95% CI)/p value for "bin animal" and "bin end": 3.3/(2.65, 4.13)/p < 10e-24; 1.67/(1.40, 1.98)/p<1e-9. (C-G) Single-animal GLMs. *p < 0.05, **p<0.01, ***p<0.001, ****p<0.0001.

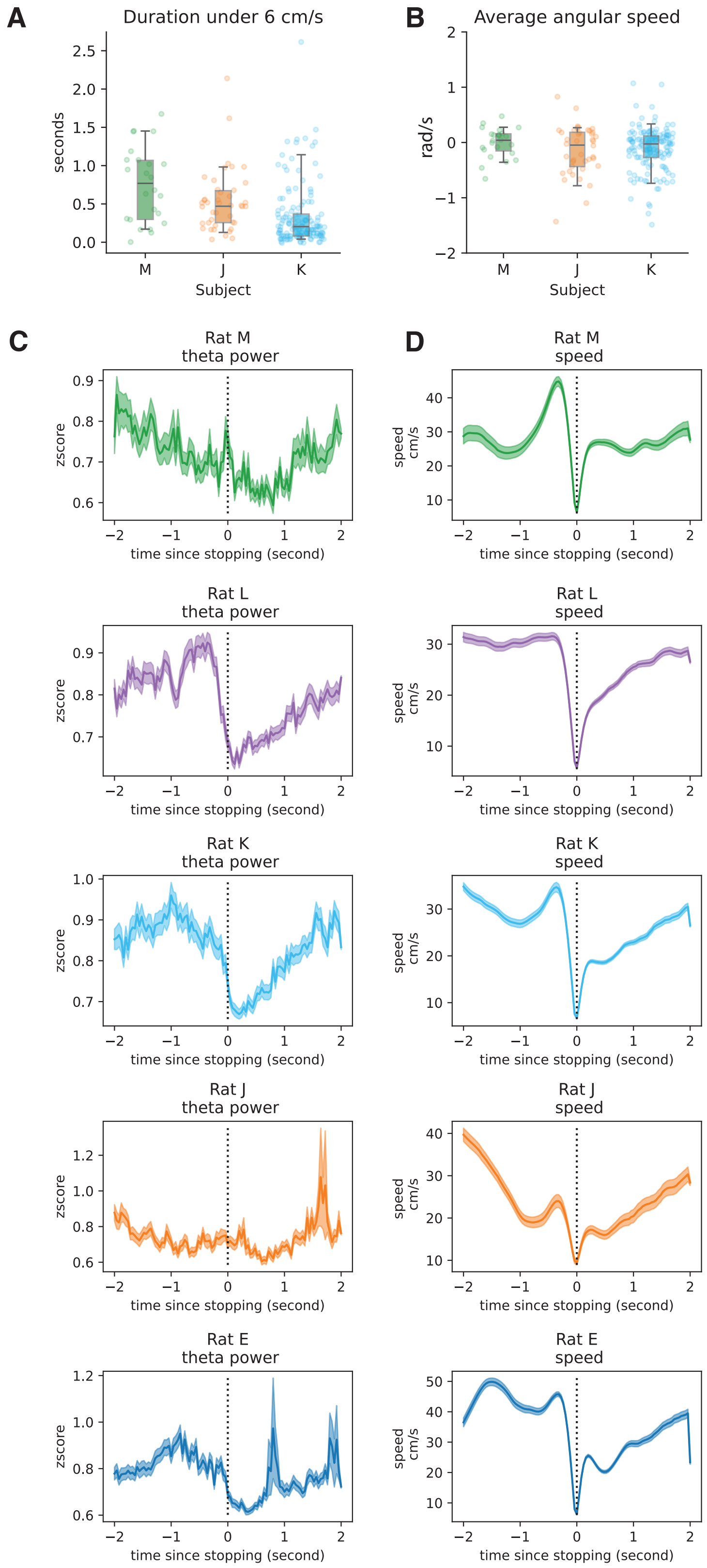


**Figure S4. Speed profile and theta power during change-of-mind.**

A. Length of periods where 2D head speed was under 6cm/s. Data shown across 3 subjects where we combined gyroscope, accelerometer data and 28Hz camera data to produce a high precision estimate of animal location and speed. Each data point comes from a COM event. Boxplots show the mean and 5$^{th}$ and 95$^{th}$ quantile of data. On average, animals were moving slowly for short periods.

B. Average angular speed from 1 second pre-stopping to 2 seconds post-stopping. Each data point comes from a COM event. Boxplots show the mean and 5$^{th}$ and 95$^{th}$ quantile of data.

C. Z-scored theta power around stopping in 5 subjects. The mean trace across all COM events is plotted in solid line and standard deviation is plotted in the shaded region.

D. 2D head speed derived from 28 Hz camera around stopping in 5 subjects. Note that the average speed does not decrease to 0 cm/s due to the jitter of 1-2 frames during the detection of stopping and the low acquisition rate. Mean across all COM events are plotted in solid line and the standard deviation is plotted in shaded region.

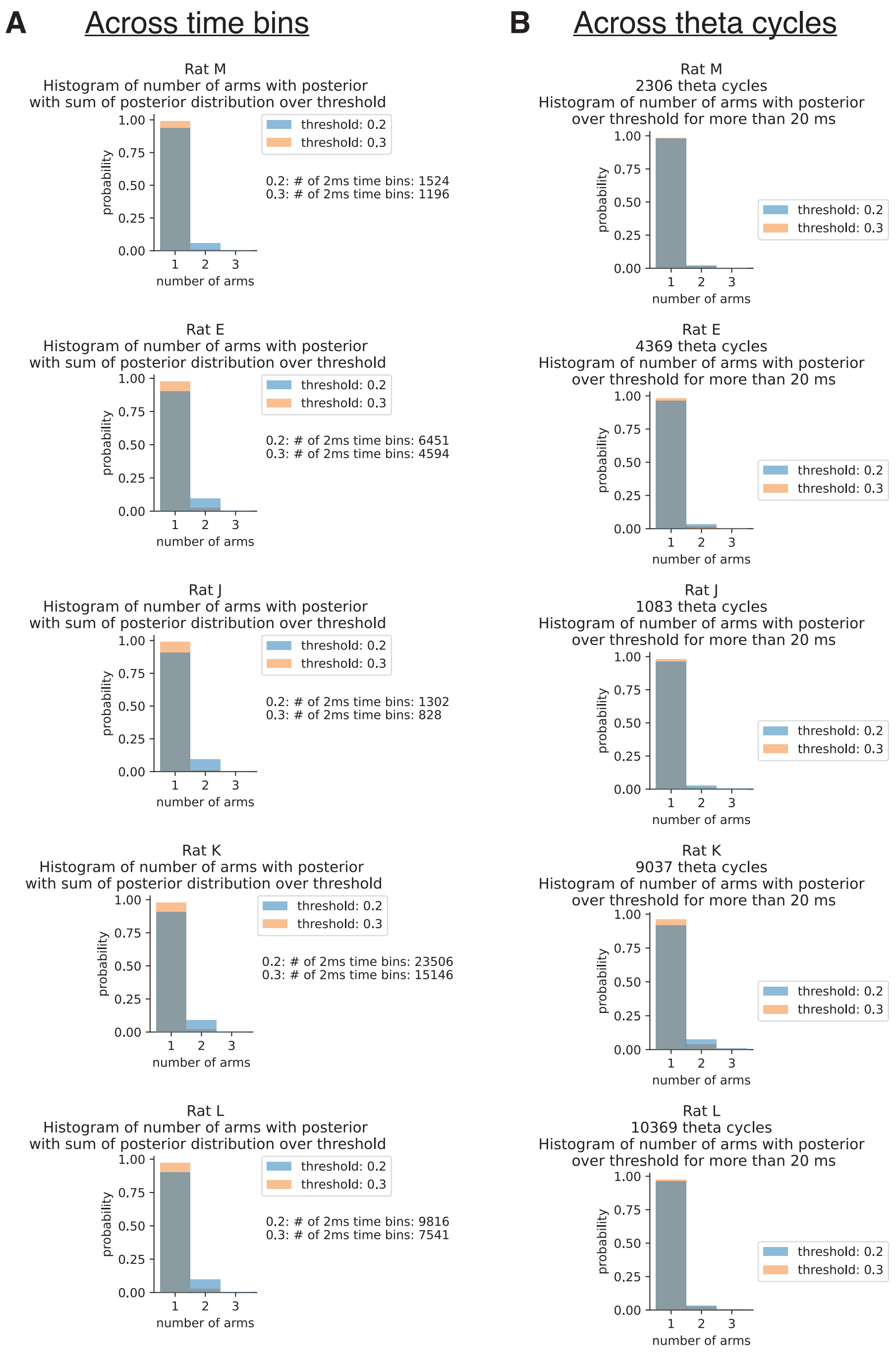


**Figure S5. Posterior of the decode is concentrated.**

Histogram of number of arms with posterior over threshold for more than 20 ms for (A) each time bin (B) theta cycle. Histograms due to a threshold of 0.2 and 0.3 are plotted in blue and orange respectively.

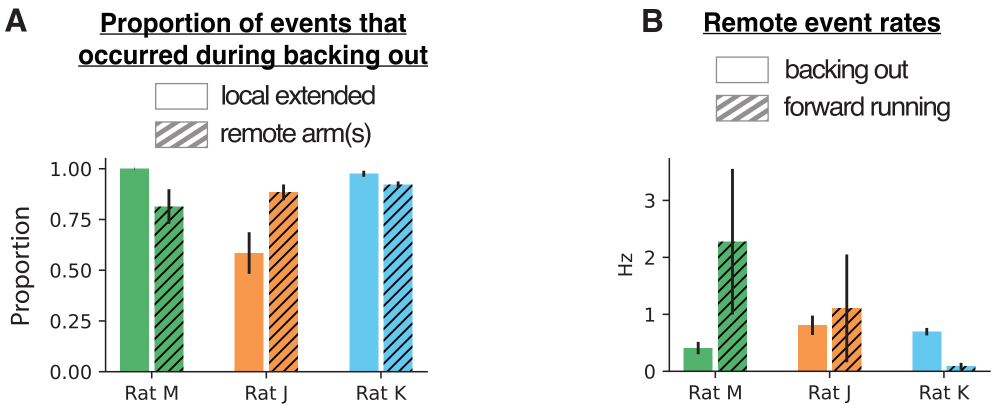


**Figure S6. Local extended or remote content on COM trials is observed both during backing out and normal running**

**A.** Proportion of events that occurred during backing out behavior. Local extended in empty bars; remote events in hatched bars. Error bars denote standard error.

**B.** Remote event rates during backing out (empty bars) or normal running (hatched bars). Error bars denote standard error.

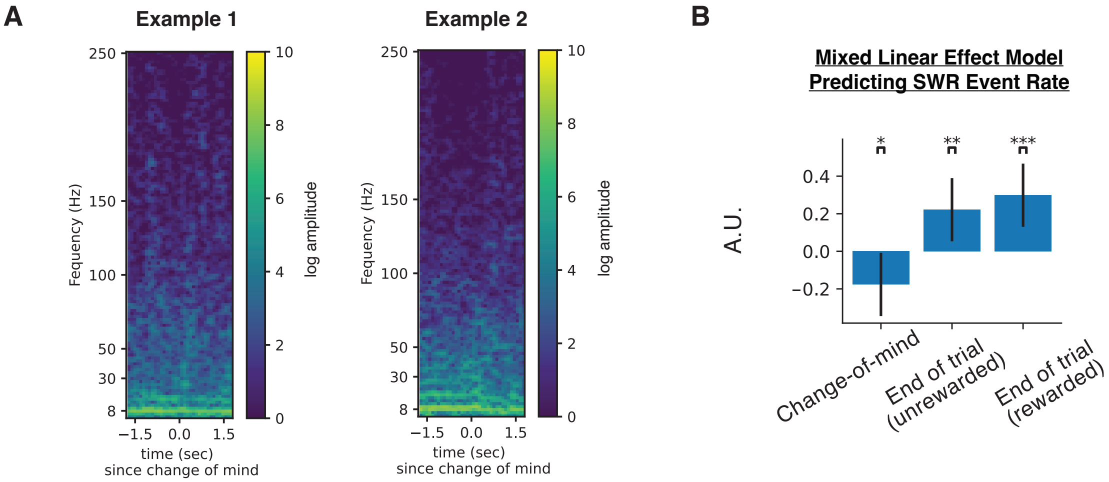


**Figure S7. Spectrogram and lack of SWR events during changes-of-mind.**

In Figure S7 – S10, we quantify the extent to which these remote representations were organized by the theta rhythm. Fig S8: phase relationship to content; Figure 9: remote content duration; Figure 10; theta power and cycle lengths.

A. LFP spectrogram during 2 seconds around COM with amplitude in logscale. Examples 1 and example 2 correspond to the COM events in Figure 2. Spectrum was calculated from one channel of each tetrode in the hippocampal CA1 cell layer with a 500 ms time window. Windows overlapped by 400 ms (100 ms step size) and then averages were computed across tetrodes.

B. Beta coefficients from predicting ripple rate per trial during COM events or at the end of a trial with rewarded or unrewarded outcomes. End of trials data are randomly sampled to match the number of COM trials. Mean estimate of the coefficients, 95% confidence interval and p values are: change of mind, -0.18, (-0.34, - 0.02), $p < 0.05$, end of trial unrewarded, 0.22, (0.06, 0.38), $p < 0.01$, end of trial rewarded, 0.30, (0.14, 0.46), $p < 0.001$.

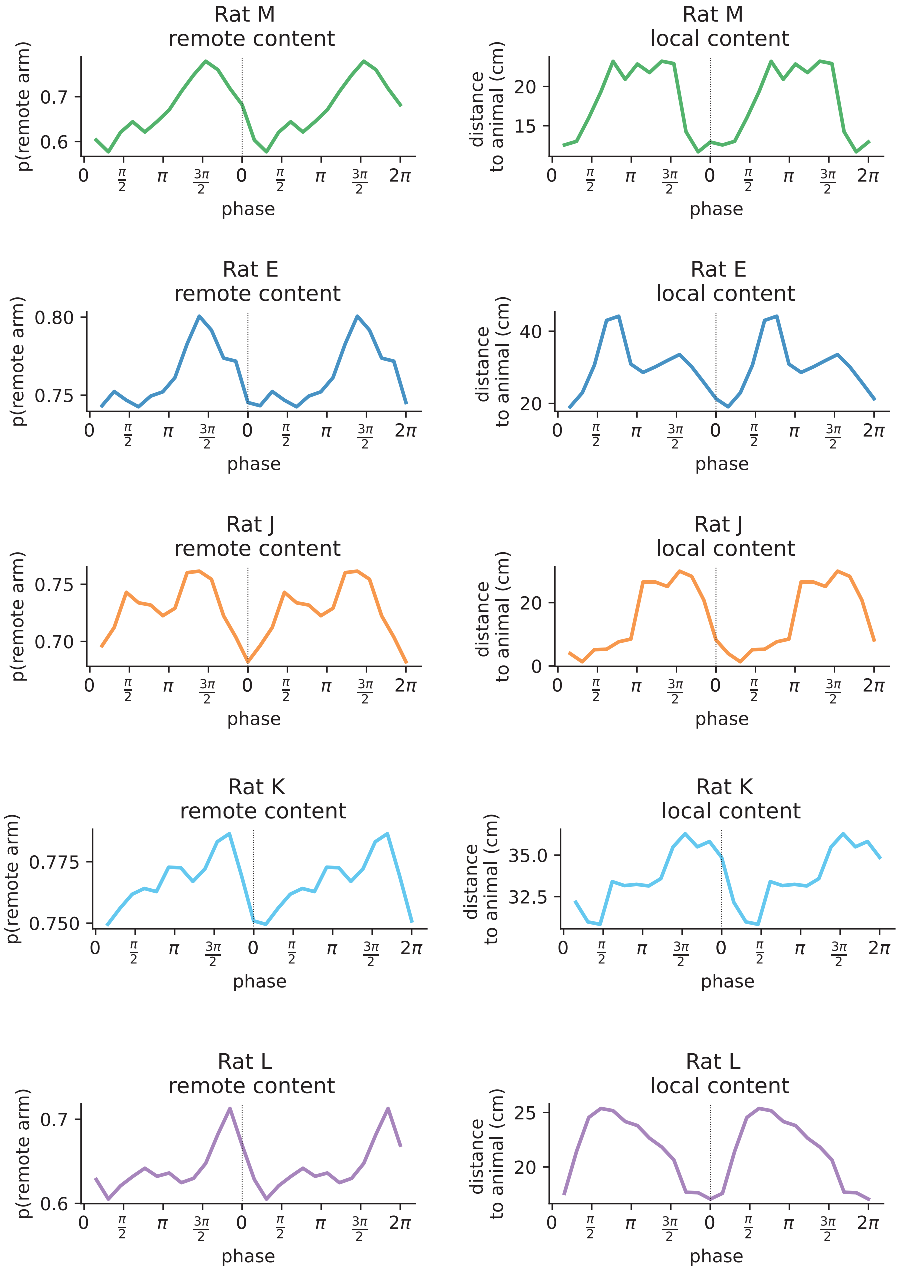


**Figure S8. Hippocampal theta phase relationship with remote content (left) and local extended content (right).** Events were more likely to occur during late theta phase. Phase 0 and $2\pi$ were defined as the peak multi-unit activities and the phases in between peaks were linear interpolated. This figure is part of the investigation into the organization of contents of alternatives by theta (Figure S7 – S10). In this figure and Figure S11 we used theta derived from multi-unit activities due to the electrodes in corpus collosum in Rat K and L have slightly sunk into the CA1 cell layer.

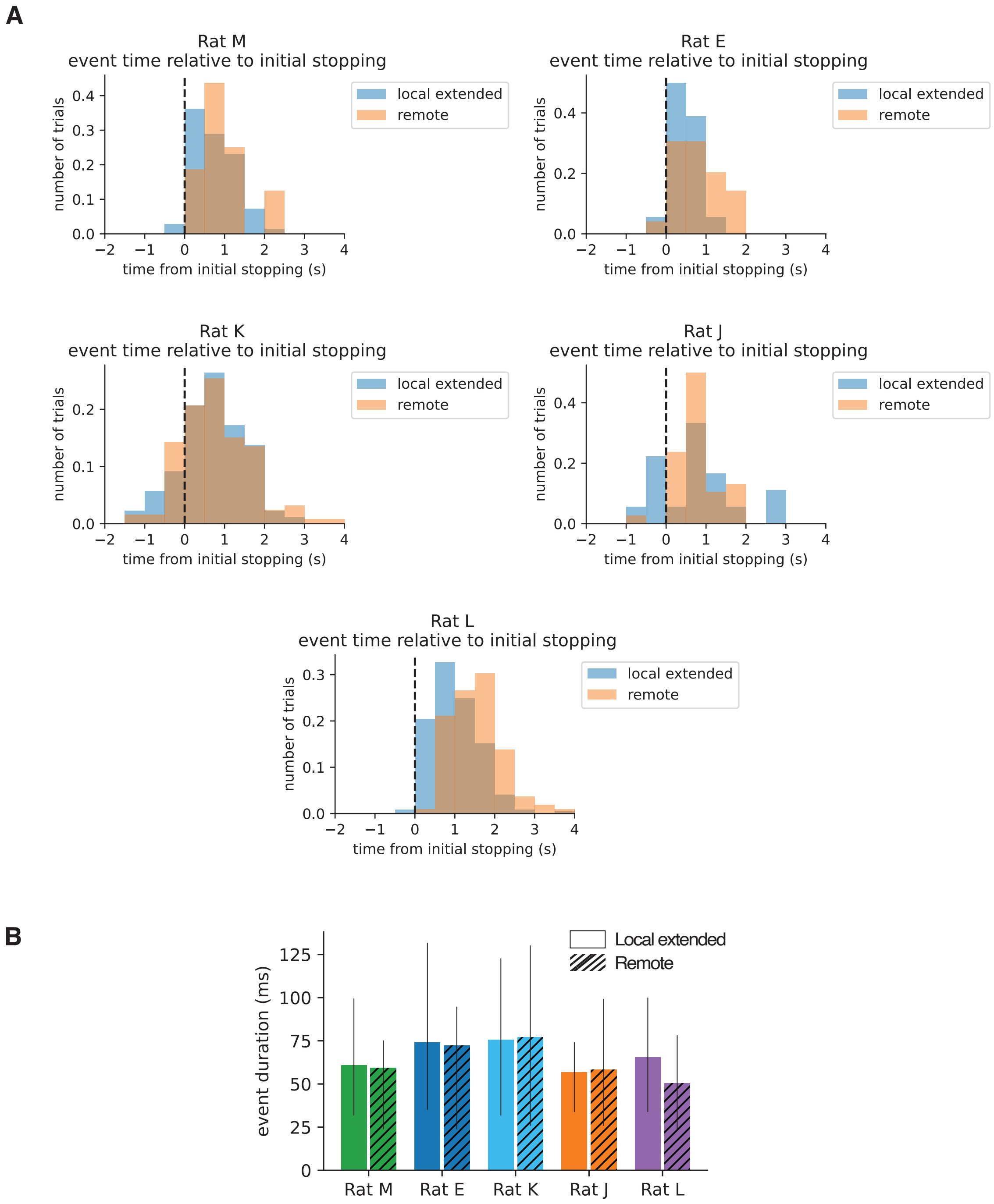


**Figure S9. Start time and end time of representations of alternatives.**

A. Start time for local extended (blue) and remote (orange) content relative to stopping in each COM event.
B. Event duration for local extended (hollow fill) and remote (hatched) contents. Events lasted for about half a theta cycle on average.

This figure is part of the investigation into the organization of contents of alternatives by theta (Figure S7 – S10).

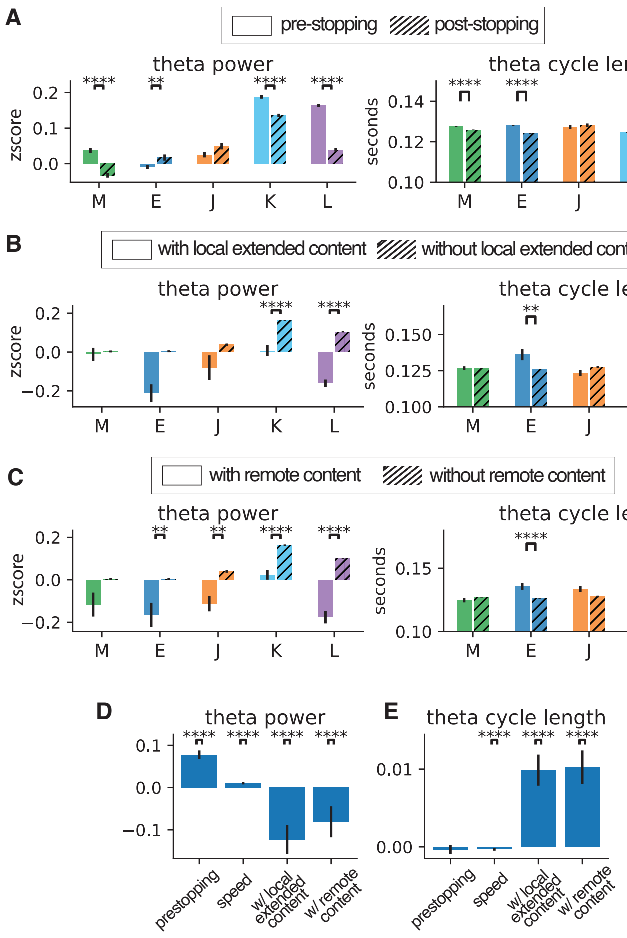


**Figure S10. Theta cycles with representations of alternatives have lower power and long durations, after controlling for movement speed.**

A. Z-scored theta power (left) and theta cycle length (right) when animal is in outer arm during a COM event, either pre-stopping (empty bars) or post-stopping (hatched bars). For panel A,B, and C, bar heights and the error bars show the mean and standard deviation respectively.
B. Z-scored theta power (left) and theta cycle length (right), for cycles with local extended content (empty bars) or those without local extended content (hatched bars).
C. Same as in B except the two categories are cycles with remote content (empty bars) or those without remote content (hatched bars).
D. and E. Beta coefficients of mixed generalized linear regression with data from all rats. Pre-stopping and a higher speed are significantly associated with higher theta power and shorter theta cycles while the existence of local extended content or remote content is significantly associated with lower theta power and longer theta cycles. $P < 0.0001$ for all significant coefficients. Bar heights and the error bars show the mean and 95% confidence interval respectively. The additive effect of local extended content and remote content on theta power is (mean, 95% CI): -0.123, (-0.154, -0.092) and -0.081, (-0.115, -0.047). The additive effect of local extended content and remote content on theta cycle duration is (mean, 95% CI): 0.010, (0.008, 0.012) and 0.010, (0.008, 0.012) second.

This figure is part of the investigation into the organization of contents of alternatives by theta (Figure S7 – S10).

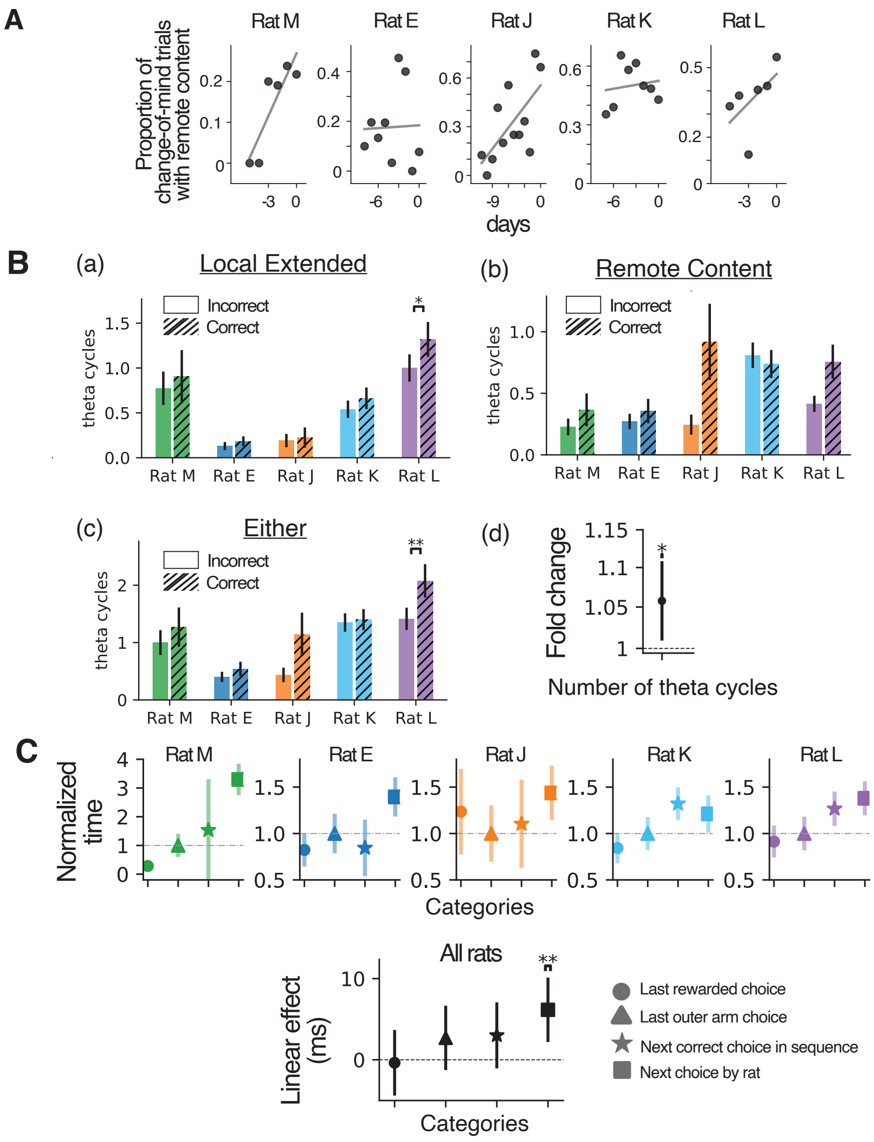


**Figure S11. Rate of remote content across days and non-binarized version of analyses as in Figure 4.**

A. COM events engaged an increasing rate of remote content across days in all five animals (Figure S11, 4A; mixed linear model mean/(CI)/p value for the effect of days 0.03 / (0.01,0.05) / $p < 0.005$)

B. Number of theta cycle with (a) local extended content (b) remote content or (c) either kind of content for COM events where the next choice is incorrect (empty bars) or correct (hatched bars). In (a)-(c), ranksum tests. (d) mixed logistic regression. Mean estimate, 95% CI, and p value of the fold change effect of number of theta cycle on correct rate is 1.06, (1.01,1.11), $p < 0.015$.

C. In Figure 4E we restrict to events with remote content >= 20ms with addition requirements (see Methods), here we consider all decode bins without any thresholding. Each 2ms decoding time bin is considered remote as long as the max posterior (MAP) is not in the same outer arm as the animal is changing its mind. In Figure 4E, if a COM event has no remote content that meets the criterion (i.e. that event has no remote content), it is skipped in analysis, but here we include all COM events. For all COM events and all remote time bins, we aggregate all the times in which the MAP is in each remote arm category. (Top) Data are normalized to the "last arm choice" category. Error bars denote normalized standard error. (Bottom) Beta coefficients from mixed linear regression pooling data from all subjects. Linear effect on addition time that a choice is represented (in milliseconds) due to immediate next choice is 6.15, 95% confidence interval: (2.38~9.93), p value < 0.002 n = 136, 281, 168, 393, 380 for rat M, E, J, K, L.

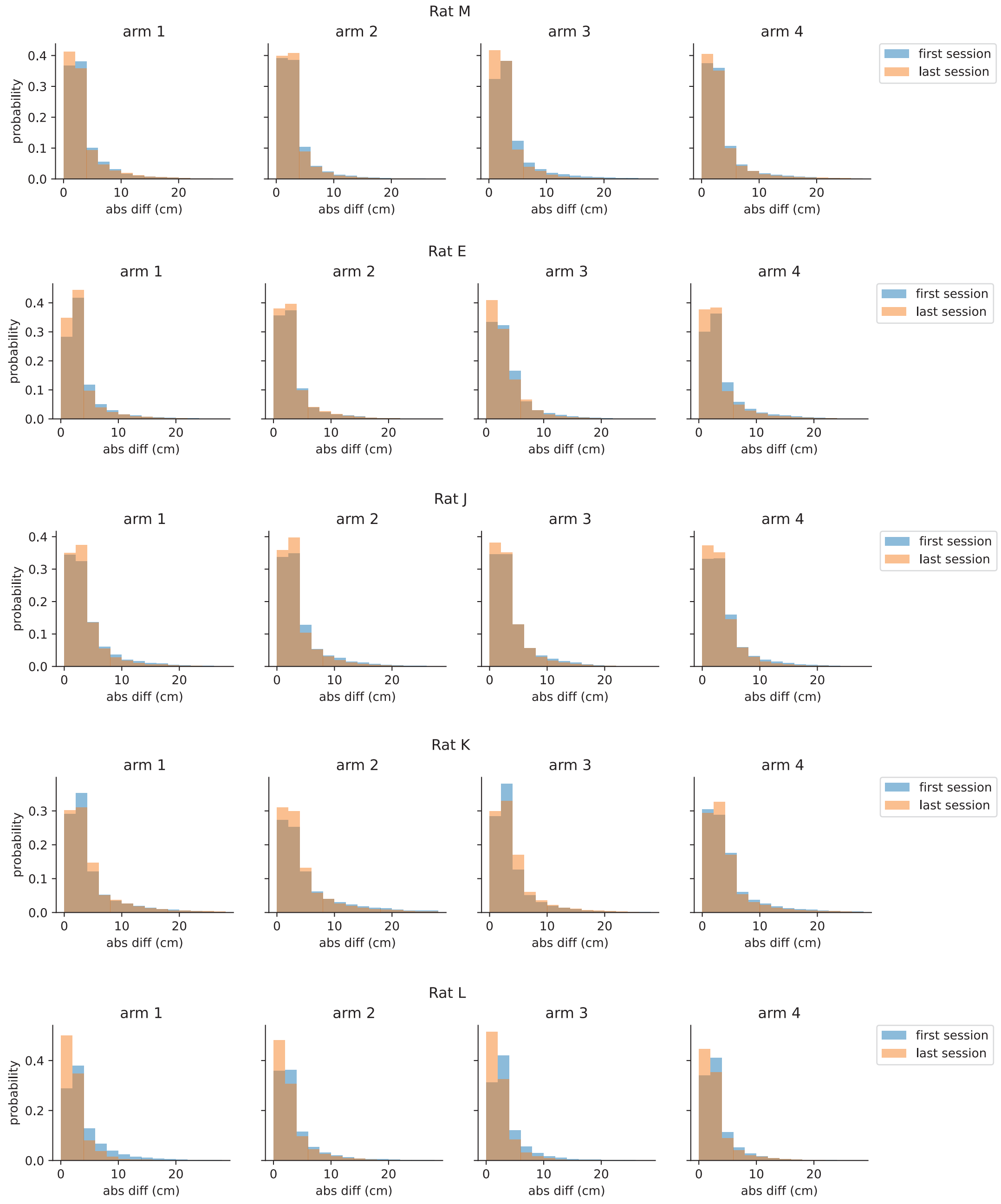


**Figure S12. Spatial decoding error is low and stable across days.**

Histograms of distance between animal head and peak posterior location of decoding. Data from the very first session in the dataset is plotted in blue, and data from the very last session is plotted in orange. Only run-time (animal head-speed >= 4cm/s and are in outer arms) are included. Data are further split by the arm the animal is in: arm 1 far left to arm 4 farther right.
